# Molecular characterization of cell types in the squid *Loligo vulgaris*

**DOI:** 10.1101/2022.03.28.485983

**Authors:** Jules Duruz, Marta Sprecher, Jenifer C. Kaldun, Alsayed Alsoudy, Heidi Tschanz-Lischer, Geert van Geest, Pamela Nicholson, Rémy Bruggmann, Simon G. Sprecher

## Abstract

Cephalopods have long been getting a lot of attention for their fascinating behavioral abilities and for the complexity of their nervous systems that set them apart from other mollusks. Because of the great evolutionary distance that separates vertebrates from mollusks, it is evident that higher cognitive features have evolved independently in this clade although they sometimes resemble cognitive functions of vertebrates. Alongside their complex behavioral abilities, cephalopods have evolved specialized cells and tissues, such as the chromatophores for camouflage or suckers to grasp prey. Gaining a better understanding of the biology of various species of cephalopods can significantly improve our knowledge of how these animals evolved and better identify the mechanisms that drive the astonishing faculties of their nervous systems. In this study, we performed single-cell transcriptomics of whole heads of *Loligo vulgaris* pre-hatchlings. We characterized the different cell types in the head of these animals and explored the expression patterns of core cell type markers by hybridization chain reaction. We were able to thoroughly describe some major components of the squid nervous that play important roles for the maintenance, development and sensory function in the nervous system of these animals.

## Introduction

Mollusks are one of the most diversified phyla in the animal kingdom: the drastic differences in body plan, morphology, and ecology between the different classes of mollusks are particularly striking. From a neuroscience perspective, these differences are also reflected by an important diversity in the organization, anatomy, shape and size of their nervous systems. Cephalopods have long piqued the interest of researchers interested in evolutionary neuroscience and are often highlighted as an ideal example of convergent evolution (Abbott et al., 1995; Young, 1965, 1985). Their large brains, relative to body size, and behavioral repertoires offer opportunities for comparative analysis of the evolution of learning and memory mechanisms. Cephalopods possess elaborate brains and are equipped with highly cognitive abilities (Darmaillacq et al., 2014; Hanlon & Messenger, 2018). They are endowed with a sophisticated nervous system that both resembles that of vertebrates in relative size (Nixon et al., 2003; Packard & Sanders, 1971) and complexity of sensory-motor systems (Hochner et al., 2006; Young, 1991). Moreover, the nervous system of cephalopods represents a distinct example of embodied organization display the highest degree of centralization (Packard, 1972) in which the central brain acts as a decision-making unit and much of the processing of motor and sensory information is performed in the peripheral nervous system. They are renowned for exhibiting strikingly flexible behavioral repertoires (Darmaillacq et al., 2014; Hanlon & Messenger, 2018; Marini et al., 2017; Mather & Dickel, 2017) such as problem-solving tasks (Fiorito et al., 1990; Richter et al., 2016), anti-predatory behaviors (Langridge et al., 2007; Staudinger et al., 2013) and social communication behaviors (Lin et al., 2007; Schnell et al., 2015).

Data obtained from fossil record suggest that cephalopods were very dominant in marine ecosystems over 300 million years ago and their biodiversity was much more important than today. The rise of bony fishes in similar ecological niches – along with other selective pressures - resulted in the extinction of many clades of cephalopods, including most of the shelled cephalopods except for the subclass Nautiloidea. However, the class cephalopod diverged into over 800 current living species with extremely diversified lifestyles and body shapes (Kröger et al., 2011). The vast majority belong to the soft-bodied subclass Coleoidea (cuttlefishes, octopuses, and squids), and a small handful belonging to the Nautiloidea (nautiluses).

The coleoid cephalopods, which have internalized or lost their shell over evolutionary time, were able to swim more easily and could exploit other niches. The continuous competition with other animals to access resources selected over hundreds of millions of years the animals that were able to develop specialized strategies that enable access to resources unavailable to others. In coleoid cephalopods, this resulted in evolved larger and highly differentiated brains, camera-type eyes as well as neutrally controlled abilities to change their skin pattern and texture for camouflage and social communication (Amodio et al., 2019; Hanlon & Messenger, 2018; Young, 1973) that enabled complex cognition, hunting and defense behaviors.

Many studies have shown that different species of cephalopods such as *Octopus vulgaris, Sepia officinalis* and *Loligo vulgaris* for example possess large and complex nervous systems and are able to perform complicated tasks. For instance, several behavioral studies have demonstrated that octopuses exhibit high flexibility in solving problems not only in their natural environment but also when faced with artificial tasks (Amodio & Fiorito, 2013; Anderson & Mather, 2010; Fiorito et al., 1990; Kuba et al., 2010; Mather & Anderson, 1999; Richter et al., 2016). The European cuttlefish, *Sepia officinalis*, has been also studied most extensively to understand the behavior (Hanlon & Messenger, 2018), learning and memory (Turchetti-Maia et al., 2017), long-term memory (Schnell et al., 2021), and the neural control of skin patterning (Hanlon, 2007; Reiter et al., 2018). These important cognitive capabilities are also reflected in their natural habitat where they use these skills to hunt, camouflage and for various social behaviors.

The European squid *Loligo vulgaris* is well-known for its economic role in the food and fishing industry but much less for its fascinating biology. Their behaviors are little studied, mostly because of practical difficulties for maintaining adult organisms in controlled laboratory conditions. However, this species is well known for its striking ability to camouflage and use its specialized iridescent cells to reflect light in different manners depending on their surroundings (Mäthger & Hanlon, 2007) and their neuroanatomy and nervous system morphology have been described in detail in histological studies (Wild et al., 2015; Young, 1976). Detailed staging of the embryonic development of the related species *Loligo pealii* was established and can be used as a reference for *Loligo vulgaris* (Arnold, 1965). While the gross anatomy of squids is well known from histology and morphological studies, we are currently lacking resolution at the molecular level to identify the genes that contribute to forming the unique cephalopod body-plan and nervous system and define its unique cell type repertoire.

Since the first complete genome sequence for *Octopus bimaculoides* was published (Albertin et al., 2015), there has been increasing interest in cephalopod biology and neuroscience. In addition, the availability of genomic sequencing for other cephalopods, such as *Euprymna scolopes* (Belcaid et al., 2019) and *Architeuthis dux* (Oegopsida) (Da Fonseca et al., 2020) as well as the genome of Nautilus (Zhang et al., 2021), opened the door to molecular comparisons between these distant clades of cephalopods with drastically different morphologies and lifestyles as a first step to understanding cephalopod evolution.

To address the lack of molecular information from squids and gain insight into their cellular diversity and gene expression, we performed single-cell transcriptomics of whole *Loligo vulgaris* heads (stage 28 based on Arnold 1965). Additionally, to validate single-cell transcriptomic analysis and explore expression of marker genes for distinct organ or cell types we used Hybridization Chain Reaction (HCR), an advanced RNA visualization technique, allowing us to spatially characterize the identified cell types. Together, these results provide new insights into the molecular and transcriptional signatures of cell types of this fascinating animal. Our detailed analysis of different aspects of the head and nervous system reveal new markers for the study of the nervous system development and function and provides key genes on interest for future studies. By analyzing the expression patterns of many cell type-specific genes, we could assess the conservation of features previously described in other cephalopods but also describe previously unknown features and modalities of cell types in this animal.

## Results

### Single cell transcriptome of the *Loligo vulgaris* head

Embryos and hatchlings at different stages were anaesthetized and sacrificed for RNA extraction and subsequent cDNA synthesis. The transcriptome was sequenced, assembled de novo, and annotated (for details see method section). To perform single-cell transcriptomics experiments, pre-hatchlings were dissected out of the eggs and were then anaesthetized. Their heads were separated and subsequently dissociated into a suspension of single cells. Single cells were captured and sequenced and the resulting barcoded sequences were clustered and analyzed. The number of cells analyzed was 20,000. Cells were filtered to keep only the ones with a gene-per-cell count of 200 to 2000 features (Median = 700 genes/cell). This filtering step resulted in a total of 19,974 cells, indicating that only 26 outliers were removed with this filtering step. Clustering was performed with Seurat using 25 principal components with a resolution value of 2. This resulted in a total of 34 cell clusters. Clusters were assigned to presumed cell type identities based on the genes differentially expressed in each individual cluster (Cl). Dimensional reduction (UMAP) was performed, and clusters were manually annotated and fitted into cell type categories based on the expression of specific markers (Fig 1B). These defined cell type categories include: muscle, connective tissue, stem cells, epidermis, papilin^+^ cells, neurons, foxD1^+^ cells, reflectin^+^ cells, ASc^+^ cells, cilia-related cells, photoreceptors, sucker and lens.

**Figure 1:**
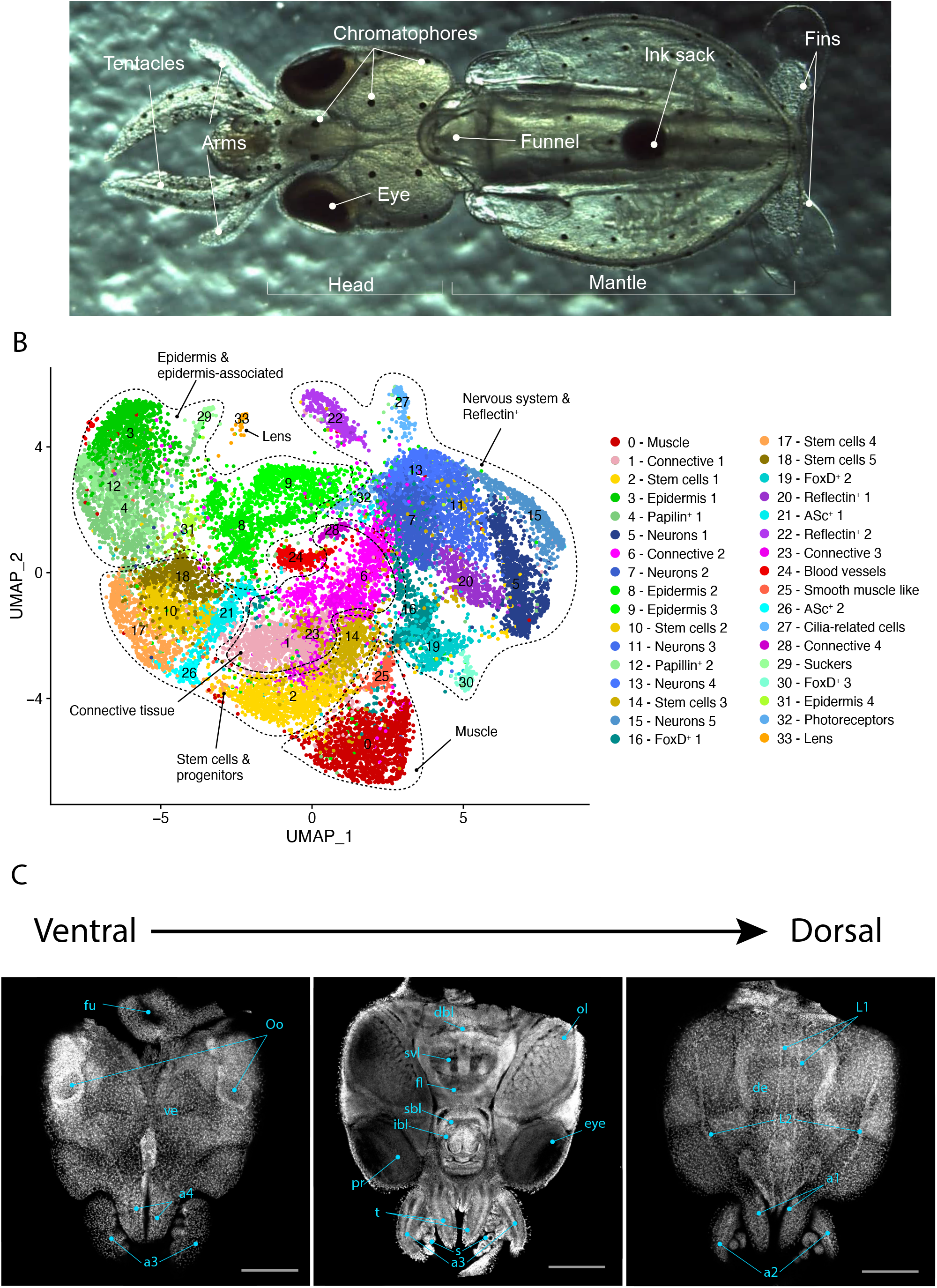
Cell atlas of the *Loligo vulgaris* head. A) Dorsal view of a recently hatched *Loligo vulgaris* with relevant external anatomical structures labeled. B) Two-dimensional UMAP representing the 34 cell clusters and their cell type characterization based on marker gene expression. These defined cell type categories include muscle, connective tissue, stem cells, epidermis, papilin^+^ cells, neurons, foxD1^+^ cells, reflectin^+^ cells, ASc^+^ cells, cilia-related cells, photoreceptors, sucker and lens. C) Successive confocal micrographs of the ventral-dorsal view of Loligo vulgaris hatchling heads stained with DAPI (nuclei). The entire ventral-dorsal surface of the head of Loligo vulgaris is labeled with relevant anatomical structures: (a) labels a1– 4=arms 1–4, (fu) funnel, (Oo) olfactory organ, (ve) ventral epidermis, (dbl) dorsal basal lobe, (svl) subvertical lobe, (fl) frontal lobe, (bl) brachial lobe, (pr) proximal segment of retinas (t) tentacle, (s) suckers, (ol) optic lobe, (de) dorsal epidermis, (L1, L2) two dorsal epidermal lines. Scale bars = 500 μM.

The identification of muscle cells was based on the expression of various genes involved in the formation of contractile fibers such as *myosin heavy chain, troponin, myosin regulatory light chain, paramyosin, tropomysin* (*Cl00*). Cl 25 expressed similar markers including *myosin catalytic light chain* and *paramyosin* but was also distinguished by the expression of the transcription factor *soxE*. Smooth muscle-like cells (*Cl26*) were characterized by the expression of the homeobox transcription factor *nkx2-5*, which has been suggested to play a crucial role in cardiac development in the human embryo (Elliott et al., 2006) and *angiopoietin-1 receptor*, involved in angiogenesis in many species. Because of the presence of these two markers, we hypothesize that these cells are in parts of the circulatory system.

Connective tissues were defined by the expression of many uncharacterized genes but could be identified by the very broad expression of many different types of collagens, which are the main component of connective tissues (*Cl01, Cl06, Cl23, Cl28*). The presence of enriched *laminin* and *fibroblast growth factor receptor* further suggests that these cells may indeed be part of connective tissues. The many uncharacterized proteins in these clusters have orthologs in other cephalopod species, suggesting that these cell types have been specialized in this clade and express a unique set of genes that may be specific to the connective tissues of cephalopods.

Cells of the epidermis were characterized based mainly on the expression of an *egf-like* epidermal growth factor (*Cl03, Cl08, Cl09*). Interestingly, *Cl08* co-expressed *tyrosinase* and *tryptophan 2,3-dioxygenase*, two members of the melanin synthesis pathway that, when knocked out, have been shown to result in depigmentation in *Doryteuthis pealeii (Crawford et al*., *2020)*. This strongly indicates that these cells are pigmented. *Cl31* was characterized by a different epithelial marker *epithelial splicing regulatory protein* (*esrp*).

Two cell clusters (*Cl04, Cl12*) highly expressed *papilin*, a gene whose function is homologous to that of a metalloproteinase in *Drosophila melanogaster (Wittmann-Liebold et al*., *1975)* and is suggested to have an important role in the formation of extracellular matrix. The exact function of this gene in other clades is however not known.

Neurons could be identified based on previously known markers such as the two members of the RNA binding elav family *elav-like* and *cugpb-elav* (*Cl05, Cl07, Cl11, Cl13, Cl15*). Known markers for the pre-synaptic machinery such as *synaptobrevin, synaptogyrin, synaptophysin* and *synapse-associated protein 1* were also broadly expressed in these clusters. Broad co-expression of genes such as *amyloid beta, tetraspanin-8, rab3* across neuronal clusters suggests that they also play an important role in the function and maintenance of the *Loligo vulgaris* nervous system.

*Cl16, Cl19* and *Cl30* were very clearly defined by the expression of the forkhead box transcription factor *foxD1*. In other phyla, this gene is involved in a variety of developmental processes. However, because the presence of this gene is not sufficient to affirm a specific cell type, we named these cells foxD^+^ cells. A large proportion of these cells co-expressed the transcription factor *ets4*. In *Cl30, foxD1* was co-expressed with a *GATA zinc-finger protein (gata-2)* and a few blood-related markers *(ferritin, hemocyte, apolipophorin)*. The possible function of these cells is discussed later in this manuscript.

*Cl20* and *Cl22* were characterized by the expression of many different types of reflectins, a group of proteins used in cephalopods to control light reflection by modulating the iridescence of the skin. Interestingly, the transcriptional repressor *scratch-2*, which is highly expressed in several neuronal clusters, was also highly expressed in reflectin^+^ cells. The expression of reflectins in neurons is further studies in later parts of the manuscript.

*Cl21* and *Cl26* were characterized by the high expression of an *achaete-scute* (*asc*) ortholog. These genes have been broadly studied for their implication in the specification of the neuronal lineage, particularly in the fruit fly *Drosophila melanogaster (Villares & Cabrera, 1987)*. These cells also expressed histone-related genes and markers of proliferation, suggesting that they are indeed undifferentiated cells. The presence of a *soxB1* homolog in these clusters also further supports that these cells may be developing neurons.

*Cl27* was characterized by the expression of different cilia-related and flagella-related markers, together with the neuronal gene *rab3* and *synaptogyrin-1*. The modality of these cells remains unknown, but the presence of cilia/flagella related genes together with neuronal markers could suggest a mechanosensory function (Fig 4).

Photoreceptors (*Cl32*) could be identified based on the expression of various members of the canonical phototransduction cascade including: *trp-like, visual arrestin, retinal binding protein, phospholipase C* and *rhodopsin*.

Cells of the lens (*Cl33*) were identified because of the expression of *s-crystallin*, a protein that in the main structural component of the tissue of the lens.

Cells of the suckers (*Cl29*) could be identified by the enriched expression of suckerin genes that have been shown to be expressed in the sucker ring teeth of squids (Kumar et al., 2016).

In the next steps of this study, we aimed to further analyze the genes expressed in cell types of particular interest, and correlate the cell types revealed by single-cell RNA sequencing with spatial analysis of gene expression, using in situ RNA visualization techniques. We aimed to further characterize the newly identified cell types and correlate them with the *Loligo vulgaris* head anatomy (Fig 1B) to gain better insights into the possible function of these cell types.

### Molecular mapping of the *Loligo vulgaris* nervous system

To assess expression patterns of newly identified neuronal markers, we performed hybridization chain reaction (HCR) to detect mRNA expression with fluorescent probes. Probes were generated for the genes *elav-like, cugbp-elav, amyloid-beta* and *tetraspanin-8*, which were all found to be expressed across all neuronal clusters (Fig 2B). HCRs for these probes confirmed their expression throughout the nervous system of *Loligo vulgaris* but different genes showed specificities in their expression domains. The expression two main members of the *elav* gene family was partly overlapping (Fig 2C). Both genes were highly expressed in the optic lobes and in the neurons of the supraesophageal and suboesophageal mass. The expression patterns show that, although they are expressed in the cortex of the same lobes within the brain and have significant areas of overlapping expression, these genes are differentially expressed in specific regions of the lobes: *elav-like* is strongly expressed in the medial part of the supraesophageal mass including neurons of the dorsal basal lobe, vertical lobe and subvertical lobe, as well as in the medial portion of the optic lobes (Fig 2C, Supplementary figure S1). *Cugbp-elav* on the other hand is expressed broadly in the lateral edges of the supraesophageal mass and in the optic lobes. Interestingly, photoreceptors of the retina appear to express *cugbp-elav* but not *elav-like* (Fig 2C, Supplementary figure S1). This suggests that these genes could serve different functions and could be defining of neuronal sub-populations based on their expression level. These different expression domains are reflected by the scRNAseq data that shows for instance that *cugbp-elav* expression level is higher than that of *elav-like* in *Cl05*, whereas the opposite can be observed in *Cl07* (Fig 2B, supplementary figure S1). Differences in the expression of these genes in the predicted cell types indicate that they may have different functions or maturation stages. The data shows that *cugpb-elav* expressing cells are present in the central and peripheral nervous system as well as in adjacent regions associated with development later in the manuscript (see Fig 3).

**Figure 2:**
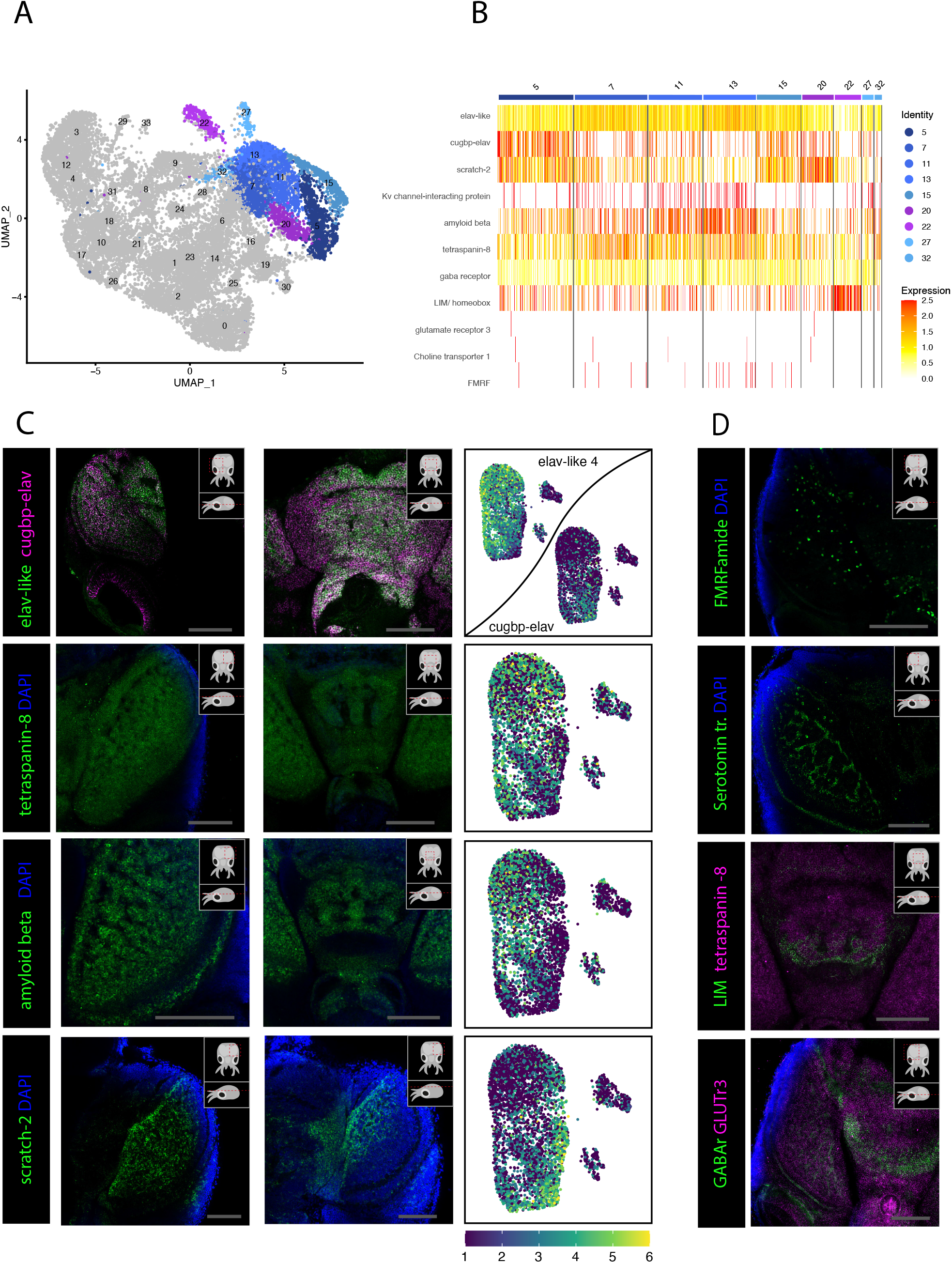
Broad makers of the *Loligo vulgaris* central nervous system. **A)** UMAP with clustered identified as neuronal or neuron-related highlighted **B)** Heatmap showing the expression of neuronal genes across each cell of the dataset. Cells are grouped by clusters. **C)** Confocal micrographs showing mRNA expression detected by hybridization chain reaction (HCR) for broad neuronal markers identified in the scRNAseq data. The right hand pictures are feature plots showing the expression of genes across the neuronal clusters in the scRNAseq data. Cartoons on the top right corner of each micrograph indicate the position of the image on a frontal view (top) and the plane of acquisition on sagittal view (bottom). Scale bars = 250 μM **D)** Confocal micrographs showing mRNA expression detected by hybridization chain reaction (HCR) for neuronal markers that were not represented in the scRNAseq data. Cartoons on the top right corner of each picture indicate the position of the image on a frontal view (top) and the plane of acquisition on sagittal view (bottom). Scale bars = 250 μM.

**Figure 3:**
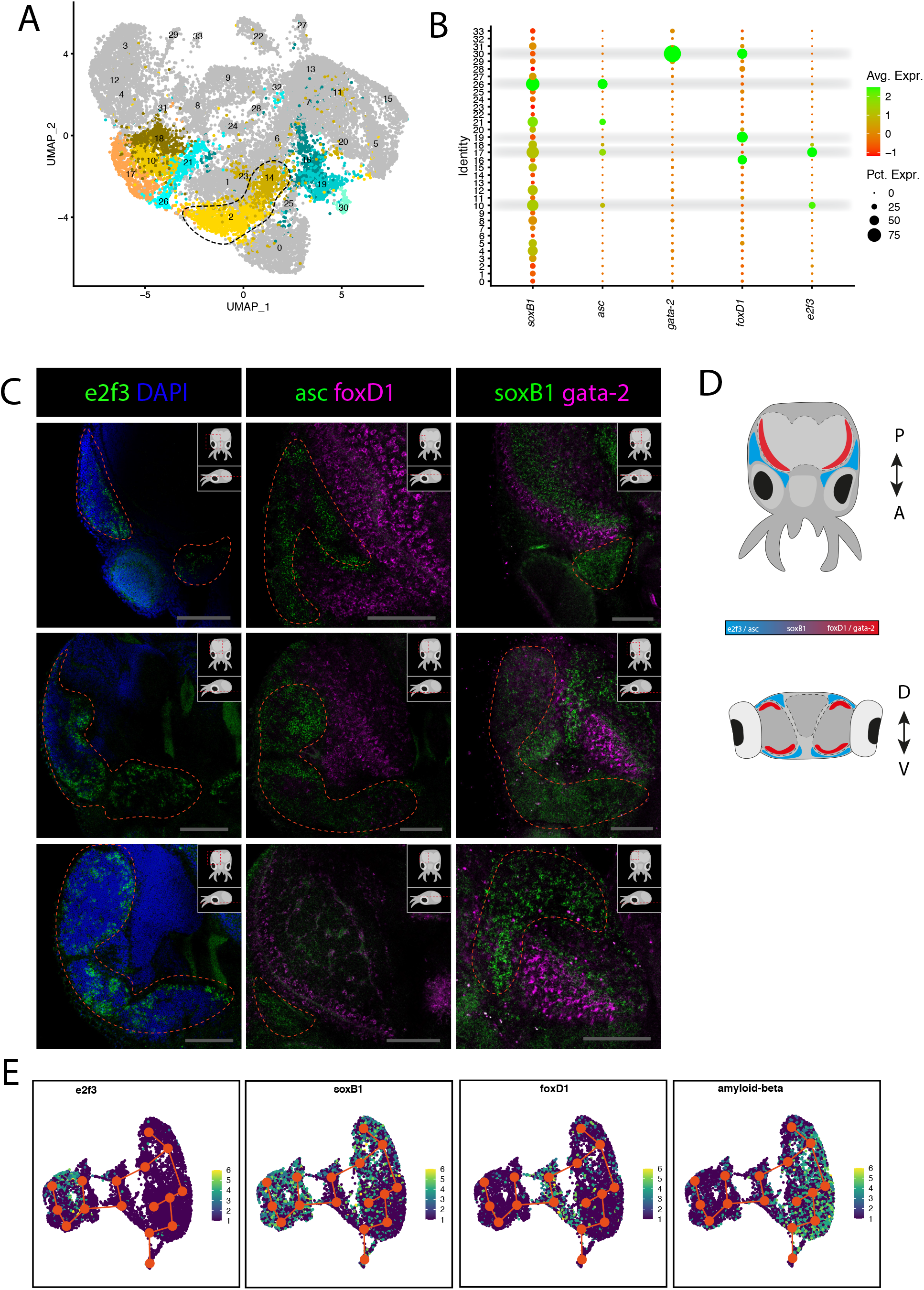
Stem cells and neurogenesis. A) UMAP with clustered identified as stem-cells, Asc^+^ or FoxD^+^ highlighted. B) Dot plot showing the average expression of marker genes and proportion of cells in each cluster that express these genes. C) Confocal micrographs showing mRNA expression detected by hybridization chain reaction (HCR) for genes involved in stem cells and neuronal development. Cartoons on the top right corner of each picture indicate the position of the image on a frontal view (top) and the plane of acquisition on sagittal view (bottom). Scale bars = 250 μM. D) Graphic representation of the gradient of gene expression in the head of *Loligo vulgaris* with dorsal view (top) and transversal view (bottom). P/A = posterior / anterior, D/V = dorsal/ventral. E) Feature plots showing the expression patterns of indicated genes across neurons and stem-cells. Cell developmental trajectories determined with slingshot are indicated in orange. The nodes correspond to the different cell clusters.

Expression of the marker *tetraspanin-8* was assessed and was observed to be present in all cells throughout the brain. The signal suggests that *tetraspanin-8* is expressed pan-neuronally. The expression pattern suggests that *tetraspanin-8* may be required during terminal differentiation of neurons or in the mature neurons in *L. vulgaris* providing an interesting candidate for the study of broad neuronal function in cephalopods. The pan-neuronal expression of *tetraspanin-8* allowed precise description of neuroanatomical regions of the *Loligo vulgaris* central nervous system (Supplementary figure S2E-G).

Similarly, the expression of *amyloid beta* was observed broadly throughout the entire brain, retina and other components of the nervous system (Fig 2C). However, at higher magnification it becomes evident that expression levels differ between cells. This is specifically striking in the optic lobe, where medially located cells of the outer granular layer and laterally located cells of the inner granular layer express lower levels of *amyloid beta* (Fig 2C, supplementary figure 2 A-D). Expression of *amyloid beta* can also be observed at lower level in the olfactory organ (supplementary figure S2C). Additionally, the scRNAseq data shows that the expression of *amyloid beta* is lower in the reflectin^+^ clusters (*Cl20, Cl22*). This indicates that this gene may be less important for these cell types that may be highly specialized (see figures 4, 5 for deeper analysis of *reflectin*^*+*^ clusters).

**Figure 4:**
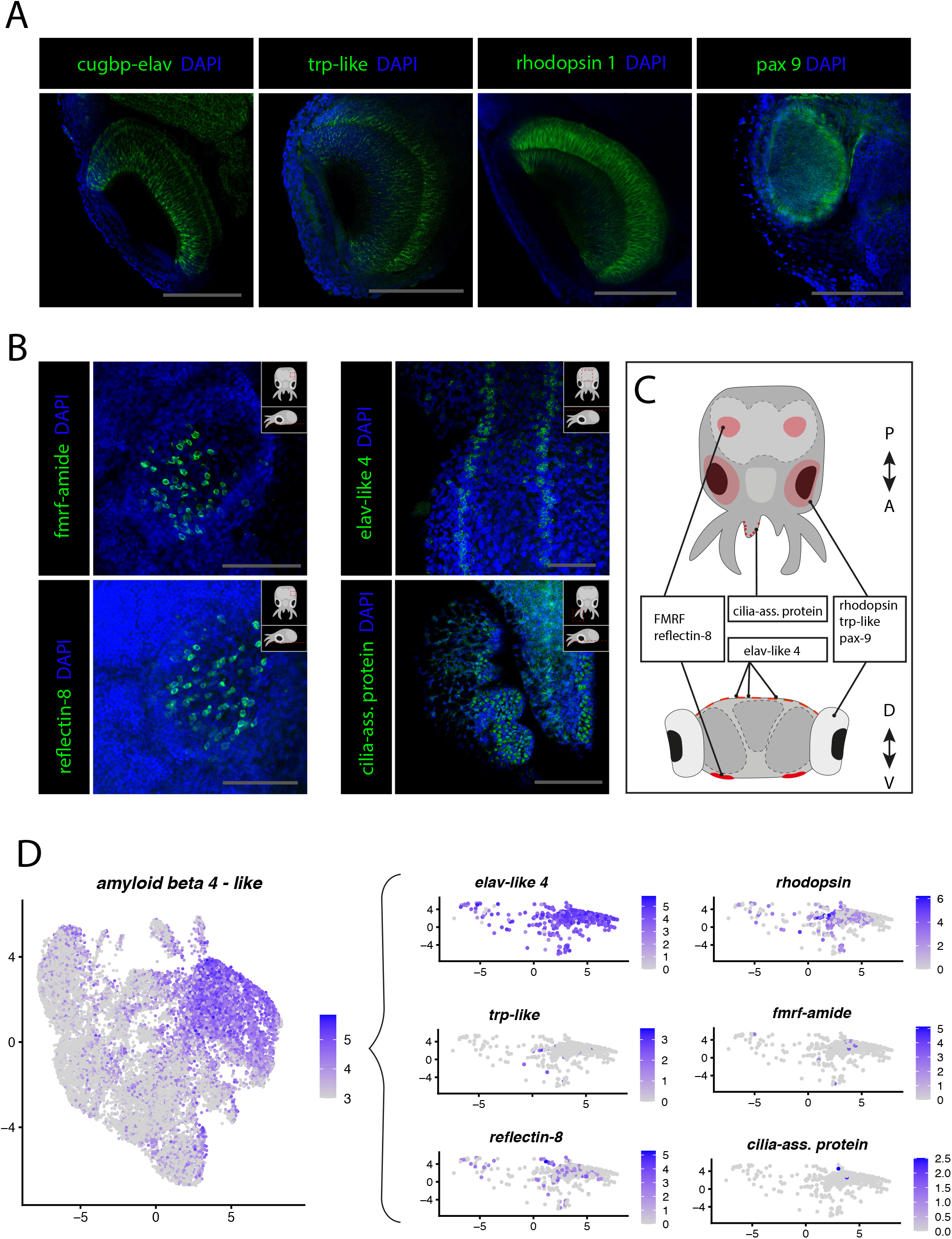
Sensory cell types. A) Confocal micrographs showing mRNA expression detected by hybridization chain reaction (HCR) for genes involved in photoreceptor function and eye development. Scale bars = 250 μM. B) Confocal micrographs showing mRNA expression detected by hybridization chain reaction (HCR) for genes involved in chemosensory and mechanosensory functions. Cartoons on the top right corner of each picture indicate the position of the image on a frontal view (top) and the plane of acquisition on sagittal view (bottom). Scale bars = 250 μM. C) Graphic representation summarizing the expression of identified genes in specific sensory organs and regions. Dorsal view (top) and transversal view (bottom). P/A = posterior / anterior, D/V = dorsal/ventral. E) Feature plots showing the cells expressing the neuronal marker amyloid-beta precursor (left). Expression of neuronal and sensory marker genes in a subset of cells amyloid-beta expressing cells above the threshold of 3.

**Figure 5:**
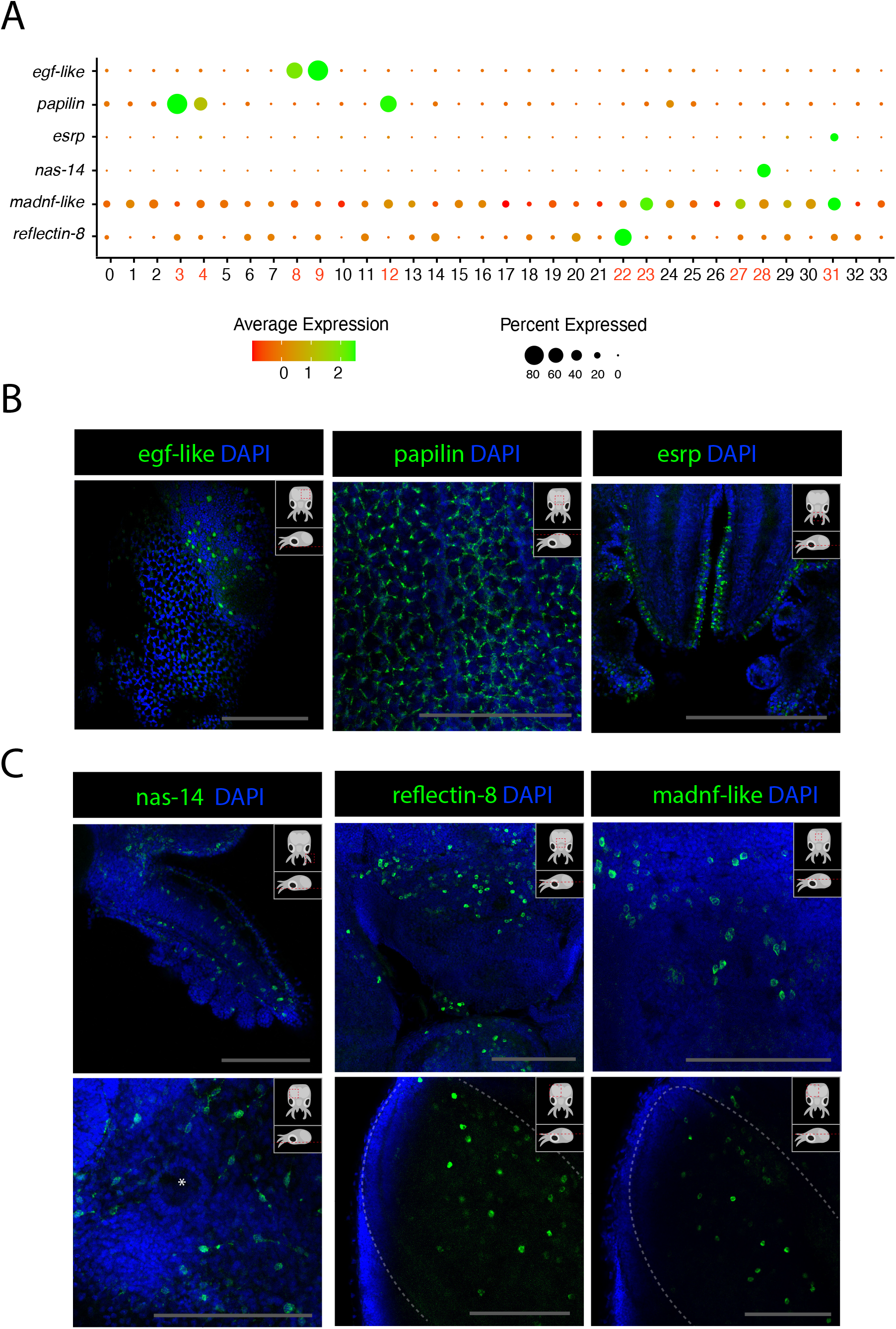
Epidermis, connective tissue and iridophores. A) Dot plot showing the average expression of marker genes of interest and proportion of cells in each cluster that express these genes. Clusters identified as epidermal, connective tissue or reflectin^+^ are labeled in red. **B)** Confocal micrographs showing mRNA expression detected by hybridization chain reaction (HCR) for genes involved in epidermis development or function. Cartoons on the top right corner of each picture indicate the position of the image on a frontal view (top) and the plane of acquisition on sagittal view (bottom). Scale bars = 250 μM. **C)** Confocal micrographs showing mRNA expression detected by hybridization chain reaction (HCR) for genes expressed in connective tissue or associated with iridophore function. Cartoons on the top right corner of each picture indicate the position of the image on a frontal view (top) and the plane of acquisition on sagittal view (bottom). Scale bars = 250 μM.

The transcriptional repressor *scratch-2* was found to be highly expressed in two neuronal clusters (*Cl05, Cl15*) as well as in a reflectin^+^ cluster (*Cl22*). HCR expression analysis showed that this gene in expressed mostly in the medial half of the optic lobes and in the supraesophageal mass in two domains lateral-ventral to the vertical lobe and anterior to the basal lobe that extends anteriorly toward the frontal lobes, as well as a stripe of cells connecting the two domains posterior to the vertical lobe (Fig 2C, Supplementary figure 3). Analysis of the expression of *scratch-2* in the scRNAseq dataset highlights that the cells highly expressing this gene are distinct from neurons expressing *elav-like, tetraspanin-8* and *amyloid-beta* but show more transcriptional similarities with *cugbp-elav* expressing cells (Fig 2C).

*Fmrf* was shown to be expressed in some cells within the nervous system but wasn ‘t as broadly expressed as previous immunohistochemical studies in cephalopods might suggest (Wollesen et al., 2010; Wollesen et al., 2012). It is expressed a region located between the dorsal basal lobe and the anterior basal lobe, surrounding the precommissural lobes, the peduncle lobe, the olfactory lobes, on the posterior part of the buccal mass and in the olfactory organ. Some cells expressing *fmrf* are also scattered throughout the optic lobes (Fig 2D, Supplementary figure 4).

The homeobox transcription factor *LIM* is expressed in the vertical lobe and subvertical lobe in a domain that separate the dorsal basal lobes from the frontal lobes. It is also expressed around the peduncle lobe and in some cells at the surface of arms and tentacles (supplementary fig S5).

Some additional genes that were not represented by the scRNAseq data were selected to relate neuronal populations to different neurotransmitters and assess whether these genes show a correlation with cell type identities or are expressed in known anatomical structures. A *serotonin transporter* gene was expressed in the nervous system and more particularly inside the optic lobes. Low expression of this gene can be expressed throughout the nervous system and indicates that the expression of the gene in the nervous system is not specific to an anatomical structure (Fig 2D). This transporter was however very highly expressed in the cornea of eyes (supplementary figure S5D-F).

Two common receptors for glutamate (*glutr3*) and GABA (*gaba-r*) were also tested with HCR. A very large proportion of neurons express *glutr3* but *gaba-r* expression was mostly restricted to the medial part of the brain and a few cells of the optic lobes (Fig 2D). More specifically *gaba-r* is expressed in the region posterior to the dorsal basal lobes, between the anterior basal lobes and frontal lobes, in the peduncle lobes and the superior buccal lobes (supplementary figure S6). *Gaba-r* appears to also be highly expressed in both neuronal and non-neural parts of the eyes but is quasi-absent from the optic lobes. Expression of *glutr3* on the other hand is observed all over the animals in neuronal tissues but also in epithelial and muscular tissues. This extremely broad expression pattern for *glutr3* indicates that this gene is likely to have other functions unrelated to neuronal communication. While it is likely that these two genes do not represent the entirety of the cells responding to either GABA or glutamate, these results suggest that glutamate signaling is likely a dominant form of neuronal communication in the *Loligo vulgaris* brain.

### Stem cells and progenitors

Several cluster markers identified from scRNAseq data suggest involvement in proliferation and possess some conserved markers of neuronal differentiation. Two clusters (*Cl02, Cl14*, Fig 3A) did not express transcription factors that can be associated with the developmental process but show high enrichment of genes involved in cellular proliferation and protein synthesis. The presence of such typical features of cycling cells could indicate that they are stem cells. In *Cl10* and *Cl17*, the transcription factor *e2f3* is expressed together with other indicators of cellular proliferation, cellular growth and nucleic acid synthesis such as histone proteins (*H3*), ribosomal proteins (*40s ribosomal protein S13, 40s ribosomal protein s19*) and DNA replication licensing factors (*mcm3, mcm4, mcm5*). This suggests that the transcription factor *e2f3* could play an important role in the identity of stem cells in *Loligo vulgaris*. To visualize the expression pattern of this gene, we performed HCR experiments for *e2f3*. The results showed that this gene is specifically expressed in anatomical structures that appear homologous to the lateral lips, which have been described in *Octopus vulgaris (Deryckere et al*., *2021)* adjacent to the nervous system and the eyes (Fig 3C-D, supplementary figure S7). This suggests the presence and proliferation of stem-cells in the lateral lips of squids, consistently with the proposed neurogenic region of *Octopus vulgaris*.

In *Cl17, Cl2*1 and *Cl26*, the transcription factor *achaete-scute homolog* (*asc*), which has a well-studied role in the regulation of nervous system development in *Drosophila melanogaster (Romani et al*., *1987)*, was expressed together with similar proliferation markers as indicated previously. This is particularly interesting to assess to possible conservation of this gene ‘s function in nervous system development. The pattern revealed by HCR showed a strong similarity with the expression pattern of *e2f3* in the lateral lips (Fig 3C-D, Supplementary figure S8), which could suggest that stem cells start to differentiate into neurons in the lateral lips before migrating to the nervous system. It remains possible that this *asc* does not play a role in nervous system development because of the lack of expression of early neuronal markers in these cells. However, frequent co-expression with *soxB1* in these clusters further supports their “neuroblast” identity because of the conserved involvement of *soxB1* genes in neuronal development in very distantly related species such as *C. elegans (Arsic et al*., *1998)* and *Homo sapiens (Alqadah et al*., *2015)*.

The expression pattern of *soxB1* was also assessed by HCR and revealed enriched expression in the lateral lips, similarly to *e2f3* and *asc*. Interestingly the pattern of expression of *soxB1* is broader than *asc* and is also expressed in most lobes of the brain (Supplementary figure S9). Although the location of these cells strongly suggests a neuronal identity, it is important to note that these cells do not co-express markers of differentiated neurons. Together the overlapping expression patterns of *e2f3, asc* and *soxB1* in domains associated to cephalopod neurogenesis and within the nervous system strongly suggest that stem cells occupy the lateral lips and then, while undergoing neuronal differentiation, migrate into the nervous system where they fully differentiate.

*Cl16, Cl19* and *Cl30* were characterized by the expression of the forkhead box transcription factor *foxD1*. FoxD genes are known to be important regulators of development, and for this reason, these cells were presumed to be differentiating cells. The fate of these cells is uncertain although recurrent co-expression of the neuronal marker *elav-like* may suggest that they are developing neurons. HCR assessment of the expression of *foxD1* showed that these cells are present in the outer layers of the optic lobes and surround the medial basal and anterior basal lobes (Fig 4C). In the optic lobes, the expression of *foxD1* can be found in the cortex of the optic lobes and in the medial and anterior basal lobes but seem most highly expressed in the inner and outer granular layers of the optic lobes (Supplementary figure S8). *Cl30* also expressed a GATA zinc finger transcription factor (*gata-2*) together with some markers typically associated to blood cells (*hemocyte protein, immunoglobulin, soma-ferritin, apolipophorin*). The expression pattern of *gata-2* is very similar to *foxD1* and also was strongly expressed in the outer layers of optic lobes (Figure 4C, Supplementary figure S9). *Gata-2* expression can also be found at the surface of the arms and in the suckers. No evidence for the expression of *gata-2* in blood-related cell types could be shown with these experiments. Developmental trajectories were calculated using slingshot (Street et al., 2018) and suggested that *foxd1*^*+*^ cells are an intermediate developmental state of neurons (Fig 3E). *Soxb1* is expressed in stem-cells and in differentiated neurons but appear to be absent in the intermediate *foxd1*-expressing stage. *E2f3* if only present in the stem cells and *amyloid beta* is only expressed in differentiated neurons as suggested by the HCR experiments.

### Cells of sensory systems

Vision is a crucial for various behaviors in open water-oriented squids such as *Loligo vulgaris*. Its eyes are very large and the optic lobes, where visual information are processed and integrated, take up a significant part of the overall volume of the squid brain. The phototransduction cascade that allows light stimuli to be transformed into an electrical signal in the neurons is well conserved among cephalopods and we were able to clearly identify photoreceptors in the scRNAseq data. Many important components of the photostransduction cascade are highly expressed in *Cl32* such as *trp-like, retinochrome, arrestin, phospholipase C* and *rhodopsin*. To verify the expression of members of the phototransduction cascade in the photoreceptors, HCR probes for *trp-like* and *rhodopsin* were selected and tested. Both genes showed striking specificity and were expressed in the photoreceptors of the retina (Fig 4A, Supplementary figure 10).

The transcription factor *pax1-like* (orthologous to human *pax9*) was also assessed as part as a screen of transcription factors of particular interest and was shown to be specifically expressed in the eye (Fig 4A). This may indicate an involvement of this gene in eye development. The expression of *pax1-like* in photoreceptors was however not concurred by the scRNAseq data analysis where co-expression with other eye-associated marker could not be shown, possibly due to the low expression level of *pax1-like*.

The olfactory organs of *Loligo vulgaris* are located on the ventral surface of the head (see Fig 1B). The genes involved in the function of these sensory organs are unknown. Therefore, lack of identifiable markers in the scRNAseq did not allow characterization of chemosensory cells of the olfactory organ. However, through screening of some cluster markers on interest, we could observe that the olfactory organ strongly expresses FMRF-related neuropeptides (*FMRFamide*). Interestingly, the pattern of expression of *FMRF amide* in the olfactory organ (Fig 4B, Supplementary figure S4) strongly resembles that of *reflectin-8* (Fig 4B, Supplementary figure S11), a gene typically involved in the function of iridophores, which are a specialized subclass of chromatophores. FMRFamide has been previously shown to be involved in the control of chromatophores in squids (Loi & Tublitz, 2000; Mobley et al., 2008; Satpute-Krishnan et al., 2006; Sweedler et al., 2000). Together, these results, in accordance with previous literature, further suggest that the modulation of chromatophores and iridophores in these animals are not only controlled in response to visual stimulation but possibly also by chemical cues.

Additionally, we observed expression of *elav-like* in neurons of the epidermal line, a mechanosensory structure that has been proposed to have a similar functions to the lateral lines of fish (Fig 4B, Supplementary figure S1) (Budelmann & Bleckmann, 1988). These stripes were suggested to be mechanosensory cells, used to sense water currents in the cuttlefish *Sepia officinalis* as well as in the squid *Lolliguncula brevis* (Komak et al., 2005). The presence of this broad neuronal marker in this structure in *Loligo vulgaris* further informs on their potential mechanosensory function, which is likely conserved in other cephalopods.

We assessed the expression of *cilia-associated protein*, which was a marker of *Cl27*. We observed that this gene is highly expressed at the surface of the arms (Fig 4B, Supplementary figure S12). Because of the co-expression of neuronal markers and additional cilia/flagella markers in this cluster we propose that these cells could be specific mechanoreceptor in the arms of *Loligo vulgaris*.

To further confirm the neuronal identity of these proposed sensory marker, we extracted a subset of neurons from our scRNAseq dataset that highly expressed the neuronal marker *amyloid beta* (expression level > 3). We then plotted the sensory markers in this subset of cells and could show that at least a subset of these neurons also expresses sensory markers. This enabled us to confirm that the sensory cells identified in this study based on the expression of specific markers are neurons.

### Epidermis and chromatophores

The epidermis of cephalopods is very peculiar and is covered with pigmented cells organized in precise patterns. Additionally, the cephalopod-specific chromatophores are colored or iridescent cells have been suggested to be modulated by the nervous system (Wardill et al., 2012).

*Cl03, Cl08, Cl09* and *Cl31* could be identified as epidermal cells from the scRNAseq data because of the consistent expression of the epidermal growth factor EGF-like. *Egf-like* was expressed in Cl03 and co-expressed some genes involved in metabolism. In *Cl08* and *Cl09, egf-like* was co-expressed with feeding circuit activating peptides and an aminopeptidase (*xaa-pro aminopeptidase)*. In *Cl08, tyrosinase* was also expressed, which suggests that those cells may be pigmented cells due to the involvement of tyrosinase in the melanin synthesis pathway. HCR experiments showed that *egf-like* is expressed in several cells across the epidermis of the animal (Figure 5B, Supplementary figure 13).

*Cl31* was characterized as epithelial due to the expression of an *epithelial splicing regulatory protein* (*esrp*). HCR for this gene revealed that it is expressed specifically on the surface of the arms, on the epithelium covering the surface of the eyes, and in cells of the epidermis, similarly to *egf-like. Esrp* is also expressed at a lower level across the surface of the epidermis (Fig 5B, supplementary figure 12).

*Cl03, Cl04* and *Cl12* showed specific enrichment for *papilin*. In *Drosophila melanogaster* studies on the role of *papilin* showed that this protein is a secreted extracellular protein that acts as a metalloproteinase and is important for embryonic development (Kramerova et al., 2000). In *Loligo vulgaris* as well as other cephalopods, the presence of this protein is documented but its function is unknown. To locate these cell types, we performed HCR experiments *papilin* and observed that it is expressed in a very particular set cells of the skin that form characteristic rings around larger cells of the epidermis (Fig 5B). Although *papilin* is expressed everywhere on the epidermis of the head of *Loligo vulgaris*, it is absent from the surface of the olfactory organ (Supplementary figure 13). This suggests that the layering of the olfactory organ is composed of different cell types than the rest of the epidermis.

*Cl28* was characterized by the expression of a zinc metalloproteinase *nas 14-like* of unknown function. Due the important co-expression of collagen-related genes, myosins and actins, these cells were proposed to be connective tissue. HCR experiments showed expression of this gene throughout the body in large cells with extended processes (Fig 5C). The branched morphology of these cells is consistent with that of fibroblasts. *Nas-14* is expressed predominantly in the ventral epidermis but is also observed in the dorsal epidermis, the arms and tentacles and is heavily expressed on the surface of the funnel (Supplementary figure 14).

We analyzed the expression of *reflectin-8*, one of the many reflectins expressed in *Cl20* and *Cl22*. However, in addition to having been shown to be also expressed in the olfactory organ (see Fig 4), HCR for this reflectin showed broad expression at the surface of the epidermis but also inside the optic lobes and around the anterior basal lobes, superior buccal lobes and peduncle lobes (Fig 5C, Supplementary figure S11). Modulation of chromatophores and iridophores by the nervous system was previously documented (Demski, 1992). The expression pattern of *reflectin-8* shown in this study suggests that the involvement of this protein may go beyond its ability to modulate light reflection, and could play a role inside the nervous system itself.

A *mesencephalic astrocyte-derived neurotrophic factor* (*madnf*) was expressed in *Cl23, Cl27* and *Cl31* that were characterized as connective tissue, mechanosensory cells and epidermis respectively. HCR experiments for this gene surprisingly showed a very similar pattern of expression as *reflectin-8*. The results showed that this gene is expressed throughout the nervous system, including in the medial part of the optic lobes (Supplementary figure 15). It is possible that these cells, which do not express neuronal markers, are part of the “connective” tissue of the brain and may be a type of glial cells further experiments would be necessary to assess that claim.

## Discussion

In this study we provide an overview of the cell types present in the head of the *Loligo vulgaris* hatchling by combining two recently developed techniques (HCR, scRNAseq). This process allowed us to describe the nervous system in detail and identify subtypes of neurons. We were able to identify broad neuronal markers that are expressed throughout the nervous system but do show different levels of expression in predicted neuronal cell types. These newly identified neuronal markers may provide new avenues to study the broad characteristics of the unique nervous systems of cephalopods. Particularly, the genes *tetraspanin-8* and *amyloid-beta* show strikingly elevated expression in most neurons. An protein orthologous to amyloid-beta in the squid *Doryteuthis pealeii* has been investigated and was suggested to have a conserved role in cargo transport along the so-called giant axon of this species (Satpute-Krishnan et al., 2006). Our results indicate that this proposed function could be extended not just to giant axons but to most neurons including sensory neurons.

We identified key genes involved in stem-cells and development. Namely, we were able to establish overlap of expression that reflect different states of neuronal differentiation at the transcriptional level but also their migration to the nervous system. We could show that stem cells were defined by the expression of the transcription factor *e2f3* and were located in the lateral lips of the animal. Overlapping expression of *ASc*^*+*^ cells in the lateral lips together with *e2f3* indicate that these cells could represent an early stage of cellular differentiation. Co-expression of *soxB1* and *asc* in the ASc^+^ clusters predicted that they may be expressed in the same structures. However, the expression pattern of *soxB1* revealed that this gene, while also present in the lateral lips, is broadly expressed inside the nervous system. This could reflect different degrees of differentiation in these cells, where *soxB1* if present in the later stages of development during migration of these cells from the lateral lips. Given the broad presence of these cells inside nervous tissue, it is likely that these cells are neuronal. In addition, we provide valuable information regarding the different sensory cells in *Loligo vulgaris* and could identify photoreceptors and additionally could describe newly identified markers of the olfactory organ and mechanosensory cells. Finally, we further characterize the epidermis of *Loligo vulgaris* and could show with scRNAseq and HCR that the implication of the nervous system in the modulation of the epidermis is visible by the expression of the typical iridophore marker *reflectin-8* not just in the epidermis but also in the nervous system.

Together, this work provides new insights into the cell type of cephalopods. By applying single-cell RNA sequencing in a member of this fascinating clade of mollusks and by correlating this data with high-resolution mRNA imaging techniques, we can further understand the main molecular features that are defining the cell types of *Loligo vulgaris*. The analysis of the expression of several selected genes of interest can provide insight into of their function that can be starting points in future investigations of cephalopod cell types. Further studies of the nervous system of squids and the different cell types that compose them could help fill important gaps in our understanding of the evolution of the nervous system and could provide insights into the way brains complexified independently in very distant clades of animals. By comparing the molecular features of neuronal cell types of cephalopods with other species, we could better understand the evolutionary history of the brains of cephalopods and distinguish what confers them such astonishing capabilities.

## Material and methods

### *Loligo vulgaris* embryo collection and maintenance

The egg sacks were collected and were immersed in a seawater tank at 12°C, in which they were hung on strings so that they do not touch the bottom. Animals were kept until pre-hatchling stage. The stage of the animals was assessed based on the description of embryonic stages in *Loligo paelii* (Arnold 1965). As the temperature was lower in order to slow down development, the stage 28 animals were determined based on the presence of chromatophores but lack of visible ink sac and the size of the yolk sack approximately equivalent to the size of the head (Arnold 1965).

### RNA extraction

10 embryos were collected into QIAzol Lysis Reagent (Qiagen, Cat. No. / ID: 79306), homogenized and centrifuged. RNA was isolated with chloroform and precipitated with Isopropanol at -20°C. After washing with Ethanol, the RNA was resolved in DEPC-treated water.

### Transcriptome assembly

The quantity and quality of the extracted RNA was assessed using a Thermo Fisher Scientific fluorometer (Qubit 4.0) with the Qubit RNA BR Assay Kit (*Thermo Fisher Scientific, Q10211*) and an Advanced Analytical Fragment Analyzer System using a Fragment Analyzer RNA Kit (*Agilent, DNF-471*), respectively. The RNA was also tested by spectrophotometry using a *Denovix DS-11 FX* Spectrophotometer / Fluorometer to assess the purity of the RNA. Once all quality control tests confirmed high quality RNA, the “Procedure & Checklist – Iso-Seq Express Template Preparation for Sequel and Sequel II Systems” was followed (*PN 101-763-800 Version 02 (October 2019)*). Two libraries were created from the same total RNA input (300 ng): one targeting 2KB transcripts and one targeting >3Kb transcripts. Both libraries were SMRT sequenced using a Sequel binding plate 3.0, sequel sequencing plate 3.0 with a 20h movie time on a PacBio Sequel system on their own SMRT cell 1M v3 LR. The 2Kb library was loaded at 3pM and generated 33.90 Gb and 409 ’187 ≥ Q20 Circular consensus reads (CCS). While the >3Kb library was loaded at 3pM and produced 27.04 Gb and 248 ’304 ≥Q20 CCS. All steps post RNA extraction were performed at the Next Generation Sequencing Platform, University of Bern.

### Cell dissociation for single-cell RNA sequencing

The embryos were first dissected out of the eggs and their heads here removed and put in nuclease-free phosphate buffer (PBS) on ice. The heads were then briefly centrifuged at 4°C and the supernatant was carefully removed and replaced with a papain enzyme solution at a concentration 1mg/ml in nuclease-free PBS. The tissue was then incubated for 30 minutes at room temperature (20-25°C) under continuous agitation. During the incubation period, the solution was pipetted up and down every 10 minutes to facilitate dissociation. The cell suspension was then centrifuged at 2000rpm for 5min at 4°C. The supernatant was removed and replaced by 1ml of PBS containing 0.04% bovine serum albumin (BSA) to stop the enzymatic reaction. This last step was repeated one more time to ensure removal of the enzyme. The suspension was then filtered through a 40 μm Flowmi Cell strainer (Bel-Art H13680-0040). The filtered suspension was centrifuged at 2000 rpm for 5 min at 4°C. The supernatant was discarded and cells were suspended in 100μl of PBS 0.04% BSA with added RNAse inhibitor (0.1μL/ml) and further dissociation was ensured by gently pipetting the entire volume approximately 200 times. Cell concentration and viability were assessed using a DeNovix CellDrop Automated Cell Counter with an Acridine Orange (AO) / Propidium Iodide (PI) assay.

### Single-cell capture and sequencing

Single-cell RNAseq libraries were prepared using a Chromium Single Cell 3 ’ Library & Gel Bead Kit v2 or v3 (10X Genomics), according to the manufacturer’s protocol (User Guide). Two chips were loaded with the accurate volumes calculated based on the “Cell Suspension Volume Calculator Table”. The initial single-cell suspension being estimated at >600 cells/μl, we targeted to recover a maximum 10000 cells. Once GEMs were obtained, reverse transcription and cDNA amplification steps were performed.

Sequencing was done on Illumina NovaSeq 6000 S2 flow cell generating paired-end reads. Different sequencing cycles were performed for the different reads, R1 and R2. R1, contained 10X barcodes and UMIs, in addition to an Illumina i7 index and R2 contained the transcript-specific sequences

All steps were performed at the Next Generation Sequencing platform at the university of Bern.

### Single-cell RNAseq analysis

High-quality isoforms were generated from PacBio reads using the Iso-Seq analysis pipeline of single-molecule real-time (SMRT) Link (version 9), which generates circular consensus sequencing reads and then clusters and polishes the isoforms. Cd-hit-est version 4.6.8 (Fu et al., 2012) was used to remove redundancy due to different transcript isoforms. It was run using a sequence identity threshold of 95%. Afterwards, the Coding GENome reconstruction Tool (cogent v 6.0.0 (Tseng, 2020)) was used to further reduce redundancy and reconstruct the coding genome. In order to annotate the coding genome, the redundancy removed sequences were blasted against the nr database of NCBI using blastx version 2.9.0 (NCBI Resource Coordinators 2016).

*10x Genomics* Cell Ranger version 3.0.2 (Zheng et al., 2017) was used to map the single cell RNAseq reads from 10x Genomics Chromium to the above assembled coding genome and count the number of reads overlapping with each gene to produce a count matrix for each sample. Afterwards, the R package scater v. 1.14.0 (McCarthy et al., 2017) was used to automatically filter low-quality cells (cells with low library size, low number of expressed genes and high proportion of mitochondrial reads). Additionally, the gene NP-062835.1 was removed due to its very high expression. The data was normalized based on a deconvolution method (Lun et al., 2016) integrated in the R package scran v. 1.14.5 (Lun et al. 2016). Integration analysis, dimensionality reduction (using the first 25 PCs) and clustering (using a resolution of 2) were done using the R package Seurat v. 3.1.3(Stuart et al., 2019). Upregulated genes for each cluster were identified using the function FindAllMarkers of Seurat. Additionally, differential expressed genes between conditions were identified for each cluster using the function FindMarkers.

### Hybridization chain reaction

Probes for hybdridization chain reaction *HCR* (Choi et al., 2018) were ordered and designed by *Molecular Instruments*^*®*^. Whole *Loligo vulgaris* animals (stage 28, Arnold 1965) were dissected out of their eggs and were then anaesthetized in 10% MgCl_2_ in artificial seawater. Once completely immobilized, the animals were fixed in 4% formaldehyde for 48 hours at room temperature. Samples were then washed three times with phosphate buffer (PBS) and were dehydrated and stored in 100% ethanol at -20°C. Prior to HCR experiments, samples were re-hydrated in PBS in three 5min steps.

Our HCR protocol used was adapted from the provided “generic sample in solution” protocol (https://files.molecularinstruments.com/MI-Protocol-RNAFISH-GenericSolution-Rev7.pdf, Choi et al., 2018) with the following modifications: a light proteinase K treatment was performed before pre-hybridization for 5min at a concentration of 1ul/ml in PBS at room temperature and subsequently washed three times with PBS. All volumes were reduced to 200uL per well. Incubation times for hybridization and amplification were increased to 16-20h to increase signal (inappropriate for RNA quantification). DAPI was added together with one of the 5x SSCT washes. Double HCR experiments were performed by combining B1-Alexa-488 hairpins with B2-Alexa-647 hairpins.

### Confocal imaging

Images were taken on Leica Stellaris 8 falcon or a Leica TCS Sp5 laser scanning confocal microscopes. All mosaic images were taken by automatically stitching 3×3 or 3×4 confocal image stacks with a 10% overlap. Laser power and gain were adjusted depending on the experiment to ensure optimal visualization. Images were treated with ImageJ where contrast was adjusted. All images are optimized for qualitative assessment and are therefore not appropriately acquires or treated for quantitative analysis.

## References

Abbott, N. J., Williamson, R., & Maddock, L. (1995). Cephalopod neurobiology: neuroscience studies in squid, octopus, and cuttlefish. Oxford University Press Oxford.

Albertin, C. B., Simakov, O., Mitros, T., Wang, Z. Y., Pungor, J. R., Edsinger-Gonzales, E., Brenner, S., Ragsdale, C. W., & Rokhsar, D. S. (2015). The octopus genome and the evolution of cephalopod neural and morphological novelties. Nature, 524(7564), 220–224.

Alqadah, A., Hsieh, Y. W., Vidal, B., Chang, C., Hobert, O., & Chuang, C. F. (2015). Postmitotic diversification of olfactory neuron types is mediated by differential activities of the HMG-box transcription factor SOX-2. The EMBO journal, 34(20), 2574–2589.

Amodio, P., Boeckle, M., Schnell, A. K., Ostojíc, L., Fiorito, G., & Clayton, N. S. (2019). Grow smart and die young: why did cephalopods evolve intelligence? Trends in ecology & evolution, 34(1), 45–56.

Amodio, P., & Fiorito, G. (2013). Observational and other types of learning in Octopus. In Handbook of Behavioral Neuroscience (Vol. 22, pp. 293–302). Elsevier.

Anderson, R. C., & Mather, J. A. (2010). It ‘s all in the cues: Octopuses (Enteroctopus dofleini) learn to open jars. Ferrantia, 59, 8–13.

Arnold, J. M. (1965). Normal embryonic stages of the squid, Loligo pealii (Lesueur). The biological bulletin, 128(1), 24–32.

Arsic, N., Rajic, T., Stanojcic, S., Goodfellow, P., & Stevanovic, M. (1998). Characterisation and mapping of the human SOX14 gene. Cytogenetic and Genome Research, 83(1-2), 139–146.

Belcaid, M., Casaburi, G., McAnulty, S. J., Schmidbaur, H., Suria, A. M., Moriano-Gutierrez, S., Pankey, M. S., Oakley, T. H., Kremer, N., & Koch, E. J. (2019). Symbiotic organs shaped by distinct modes of genome evolution in cephalopods. Proceedings of the National Academy of Sciences, 116(8), 3030–3035.

Budelmann, B. U., & Bleckmann, H. (1988). A lateral line analogue in cephalopods: water waves generate microphonic potentials in the epidermal head lines ofSepia andLolliguncula. Journal of Comparative Physiology A, 164(1), 1–5.

Choi, H. M., Schwarzkopf, M., Fornace, M. E., Acharya, A., Artavanis, G., Stegmaier, J., Cunha, A., & Pierce, N. A. (2018). Third-generation in situ hybridization chain reaction: multiplexed, quantitative, sensitive, versatile, robust. Development, 145(12), dev165753.

Crawford, K., Quiroz, J. F. D., Koenig, K. M., Ahuja, N., Albertin, C. B., & Rosenthal, J. J. (2020). Highly efficient knockout of a squid pigmentation gene. Current Biology, 30(17), 3484-3490. e3484.

Da Fonseca, R. R., Couto, A., Machado, A. M., Brejova, B., Albertin, C. B., Silva, F., Gardner, P., Baril, T., Hayward, A., & Campos, A. (2020). A draft genome sequence of the elusive giant squid, Architeuthis dux. GigaScience, 9(1), giz152.

Darmaillacq, A.-S., Dickel, L., & Mather, J. (2014). Cephalopod cognition. Cambridge University Press.

Demski, L. S. (1992). Chromatophore Systems in Teleosts and Cephalopods: A Levels Oriented Analysis of Convergent Systems (Part 1 of 2). Brain, behavior and evolution, 40(2-3), 141–148.

Deryckere, A., Styfhals, R., Elagoz, A. M., Maes, G. E., & Seuntjens, E. (2021). Identification of neural progenitor cells and their progeny reveals long distance migration in the developing octopus brain. Elife, 10, e69161.

Elliott, D. A., Solloway, M. J., Wise, N., Biben, C., Costa, M. W., Furtado, M. B., Lange, M., Dunwoodie, S., & Harvey, R. P. (2006). A tyrosine-rich domain within homeodomain transcription factor Nkx2-5 is an essential element in the early cardiac transcriptional regulatory machinery.

Fiorito, G., von Planta, C., & Scotto, P. (1990). Problem solving ability of Octopus vulgaris lamarck (Mollusca, Cephalopoda). Behavioral and neural biology, 53(2), 217–230.

Fu, L., Niu, B., Zhu, Z., Wu, S., & Li, W. (2012). CD-HIT: accelerated for clustering the next-generation sequencing data. Bioinformatics, 28(23), 3150–3152.

Hanlon, R. (2007). Cephalopod dynamic camouflage. Curr Biol, 17(11), R400–404. https://doi.org/10.1016/j.cub.2007.03.034

Hanlon, R. T., & Messenger, J. B. (2018). Cephalopod behaviour. Cambridge University Press.

Hochner, B., Shomrat, T., & Fiorito, G. (2006). The octopus: a model for a comparative analysis of the evolution of learning and memory mechanisms. The biological bulletin, 210(3), 308–317.

Komak, S., Boal, J. G., Dickel, L., & Budelmann, B. U. (2005). Behavioural responses of juvenile cuttlefish (Sepia officinalis) to local water movements. Marine and Freshwater Behaviour and Physiology, 38(2), 117–125.

Kramerova, I. A., Kawaguchi, N., Fessler, L. I., Nelson, R. E., Chen, Y., Kramerov, A. A., Kusche-Gullberg, M., Kramer, J. M., Ackley, B. D., & Sieron, A. L. (2000). Papilin in development; a pericellular protein with a homology to the ADAMTS metalloproteinases. Development, 127(24), 5475–5485.

Kröger, B., Vinther, J., & Fuchs, D. (2011). Cephalopod origin and evolution: a congruent picture emerging from fossils, development and molecules: extant cephalopods are younger than previously realised and were under major selection to become agile, shell-less predators. Bioessays, 33(8), 602–613.

Kuba, M. J., Byrne, R. A., & Burghardt, G. M. (2010). A new method for studying problem solving and tool use in stingrays (Potamotrygon castexi). Animal cognition, 13(3), 507–513.

Kumar, A., Kannan, S., Lescar, J., Verma, C., & Miserez, A. (2016). Squid ‘s Suckerin Proteins in Bits & Bytes. Biophysical Journal, 110(3), 341a.

Langridge, K. V., Broom, M., & Osorio, D. (2007). Selective signalling by cuttlefish to predators. Current Biology, 17(24), R1044–R1045.

Lin, P.-Y., Hinterneder, J. M., Rollor, S. R., & Birren, S. J. (2007). Non-cell-autonomous regulation of GABAergic neuron development by neurotrophins and the p75 receptor. Journal of Neuroscience, 27(47), 12787–12796.

Loi, P. K., & Tublitz, N. J. (2000). Roles of glutamate and FMRFamide-related peptides at the chromatophore neuromuscular junction in the cuttlefish, Sepia officinalis. Journal of Comparative Neurology, 420(4), 499–511.

Lun, A. T., Bach, K., & Marioni, J. C. (2016). Pooling across cells to normalize single-cell RNA sequencing data with many zero counts. Genome biology, 17(1), 1–14.

Marini, G., De Sio, F., Ponte, G., & Fiorito, G. (2017). Behavioral analysis of learning and memory in cephalopods.

Mather, J. A., & Anderson, R. C. (1999). Exploration, play and habituation in octopuses (Octopus dofleini). Journal of Comparative Psychology, 113(3), 333.

Mather, J. A., & Dickel, L. (2017). Cephalopod complex cognition. Current Opinion in Behavioral Sciences, 16, 131–137.

Mäthger, L. M., & Hanlon, R. T. (2007). Malleable skin coloration in cephalopods: selective reflectance, transmission and absorbance of light by chromatophores and iridophores. Cell and tissue research, 329(1), 179–186.

McCarthy, D. J., Campbell, K. R., Lun, A. T., & Wills, Q. F. (2017). Scater: pre-processing, quality control, normalization and visualization of single-cell RNA-seq data in R. Bioinformatics, 33(8), 1179–1186.

Mobley, A. S., Michel, W. C., & Lucero, M. T. (2008). Odorant responsiveness of squid olfactory receptor neurons. The Anatomical Record: Advances in Integrative Anatomy and Evolutionary Biology: Advances in Integrative Anatomy and Evolutionary Biology, 291(7), 763–774.

Nixon, M., Young, J. Z., & Young, J. Z. (2003). The brains and lives of cephalopods. Oxford University Press.

Packard, A. (1972). Cephalopods and fish: the limits of convergence. Biological Reviews, 47(2), 241–307.

Packard, A., & Sanders, G. D. (1971). Body patterns of Octopus vulgaris and maturation of the response to disturbance. Animal Behaviour, 19(4), 780–790.

Reiter, S., Hülsdunk, P., Woo, T., Lauterbach, M. A., Eberle, J. S., Akay, L. A., Longo, A., Meier-Credo, J., Kretschmer, F., Langer, J. D., Kaschube, M., & Laurent, G. (2018). Elucidating the control and development of skin patterning in cuttlefish. Nature, 562(7727), 361–366. https://doi.org/10.1038/s41586-018-0591-3

Richter, J. N., Hochner, B., & Kuba, M. J. (2016). Pull or push? Octopuses solve a puzzle problem. PloS one, 11(3), e0152048.

Romani, S., Campuzano, S., & Modolell, J. (1987). The achaete-scute complex is expressed in neurogenic regions of Drosophila embryos. The EMBO journal, 6(7), 2085–2092.

Satpute-Krishnan, P., DeGiorgis, J. A., Conley, M. P., Jang, M., & Bearer, E. L. (2006). A peptide zipcode sufficient for anterograde transport within amyloid precursor protein. Proceedings of the National Academy of Sciences, 103(44), 16532–16537.

Schnell, A. K., Clayton, N. S., Hanlon, R. T., & Jozet-Alves, C. (2021). Episodic-like memory is preserved with age in cuttlefish. Proceedings of the Royal Society B, 288(1957), 20211052.

Schnell, A. K., Smith, C. L., Hanlon, R. T., & Harcourt, R. T. (2015). Female receptivity, mating history, and familiarity influence the mating behavior of cuttlefish. Behavioral ecology and sociobiology, 69(2), 283–292.

Staudinger, M. D., Buresch, K. C., Mäthger, L. M., Fry, C., McAnulty, S., Ulmer, K. M., & Hanlon, R. T. (2013). Defensive responses of cuttlefish to different teleost predators. The biological bulletin, 225(3), 161–174.

Street, K., Risso, D., Fletcher, R. B., Das, D., Ngai, J., Yosef, N., Purdom, E., & Dudoit, S. (2018). Slingshot: cell lineage and pseudotime inference for single-cell transcriptomics. BMC genomics, 19(1), 1–16.

Stuart, T., Butler, A., Hoffman, P., Hafemeister, C., Papalexi, E., Mauck III, W. M., Hao, Y., Stoeckius, M., Smibert, P., & Satija, R. (2019). Comprehensive integration of single-cell data. Cell, 177(7), 1888-1902. e1821.

Sweedler, J., Li, L., Floyd, P., & Gilly, W. (2000). Mass spectrometric survey of peptides in cephalopods with an emphasis on the FMRFamide-related peptides. Journal of Experimental Biology, 203(23), 3565–3573. [Record #88 is using a reference type undefined in this output style.]

Turchetti-Maia, A., Shomrat, T., & Hochner, B. (2017). The vertical lobe of cephalopods: a brain structure ideal for exploring the mechanisms of complex forms of learning and memory. The Oxford handbook of invertebrate neurobiology. Oxford University Press, Oxford, 1–27.

Villares, R., & Cabrera, C. V. (1987). The achaete-scute gene complex of D. melanogaster: conserved domains in a subset of genes required for neurogenesis and their homology to myc. Cell, 50(3), 415–424.

Wardill, T. J., Gonzalez-Bellido, P. T., Crook, R. J., & Hanlon, R. T. (2012). Neural control of tuneable skin iridescence in squid. Proceedings of the Royal Society B: Biological Sciences, 279(1745), 4243–4252.

Wild, E., Wollesen, T., Haszprunar, G., & Heß, M. (2015). Comparative 3D microanatomy and histology of the eyes and central nervous systems in coleoid cephalopod hatchlings. Organisms Diversity & Evolution, 15(1), 37–64.

Wittmann-Liebold, B., Geissler, A., & Marzinzig, E. (1975). Studies on the primary structure of 14 proteins from the large subunit of Escherichia coli ribosomes with an improved protein sequenator and with mass spectrometry. Journal of Supramolecular Structure, 3(5-6), 426–447.

Wollesen, T., Cummins, S. F., Degnan, B. M., & Wanninger, A. (2010). FMRFamide gene and peptide expression during central nervous system development of the cephalopod mollusk, Idiosepius notoides. Evolution & development, 12(2), 113–130.

Wollesen, T., Sukhsangchan, C., Seixas, P., Nabhitabhata, J., & Wanninger, A. (2012). Analysis of neurotransmitter distribution in brain development of benthic and pelagic octopod cephalopods. Journal of Morphology, 273(7), 776–790.

Young, J. (1991). Computation in the learning system of cephalopods. The biological bulletin, 180(2), 200–208.

Young, J. Z. (1965). The Croonian Lecture, 1965-The organization of a memory system. Proceedings of the Royal Society of London. Series B. Biological Sciences, 163(992), 285–320.

Young, J. Z. (1976). The nervous system of Loligo II. Suboesophageal centres. Philosophical Transactions of the Royal Society of London. B, Biological Sciences, 274(930), 101–167.

Young, J. Z. (1985). Cephalopods and neuroscience. The biological bulletin, 168(3S), 153–158.

Young, R. E. (1973). Information feedback from photophores and ventral countershading in mid-water squid.

Zhang, Y., Mao, F., Mu, H., Huang, M., Bao, Y., Wang, L., Wong, N.-K., Xiao, S., Dai, H., & Xiang, Z. (2021). The genome of Nautilus pompilius illuminates eye evolution and biomineralization. Nature Ecology & Evolution, 5(7), 927–938.

Zheng, G. X., Terry, J. M., Belgrader, P., Ryvkin, P., Bent, Z. W., Wilson, R., Ziraldo, S. B., Wheeler, T. D., McDermott, G. P., & Zhu, J. (2017). Massively parallel digital transcriptional profiling of single cells. Nature communications, 8(1), 1–12.

