## Supplemental Figures for "Molecular characterization of cell types in the squid *Loligo vulgaris*"

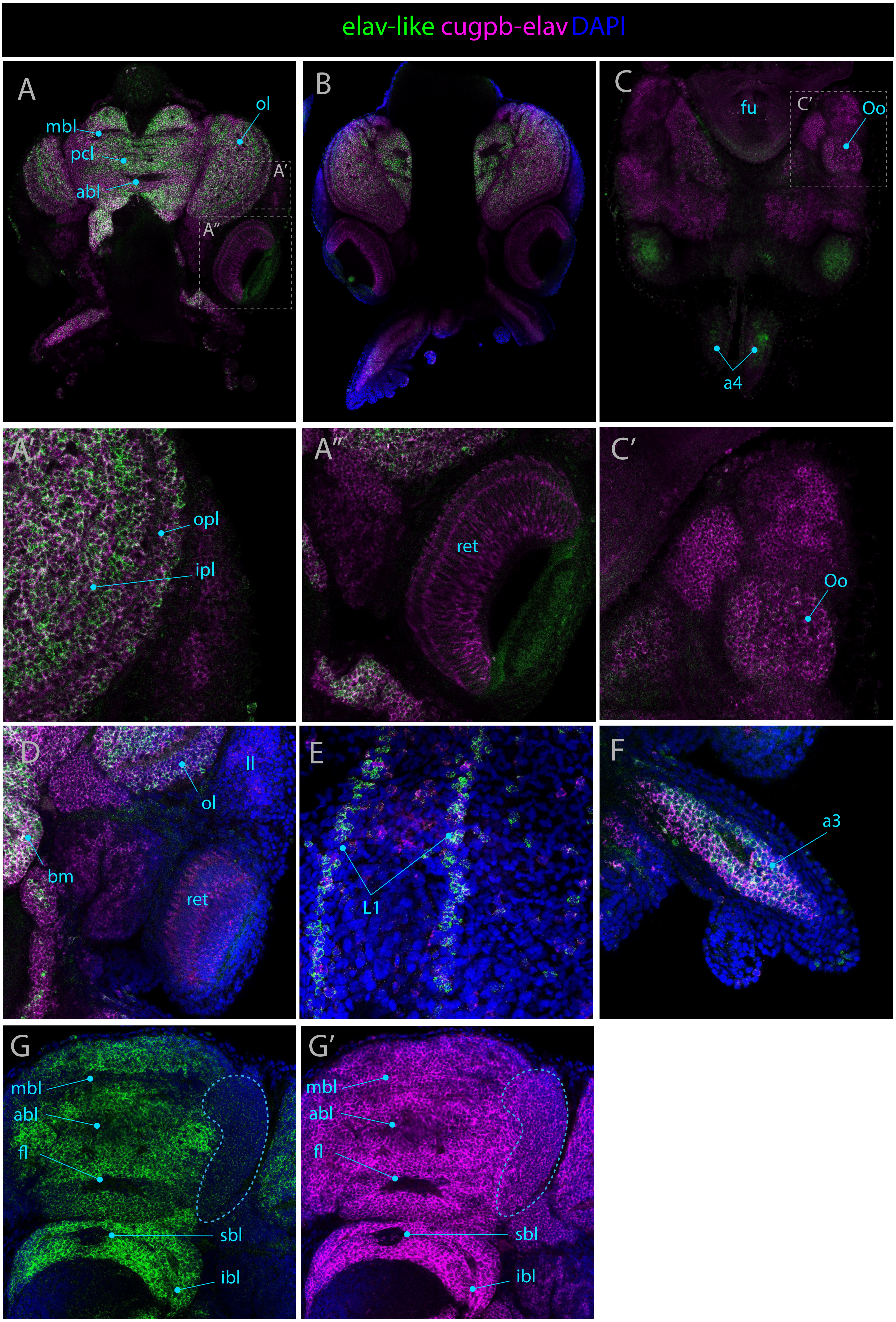

elav-like cugpb-elav DAPI
A
mbl
pcl
abl
ol
A′
A″
B
C
fu
C′
Oo
a4
A′
opl
ipl
A″
ret
C′
Oo
D
ll
ol
bm
ret
E
L1
F
a3
G
mbl
abl
fl
sbl
ibl
G′
mbl
abl
fl
sbl
ibl

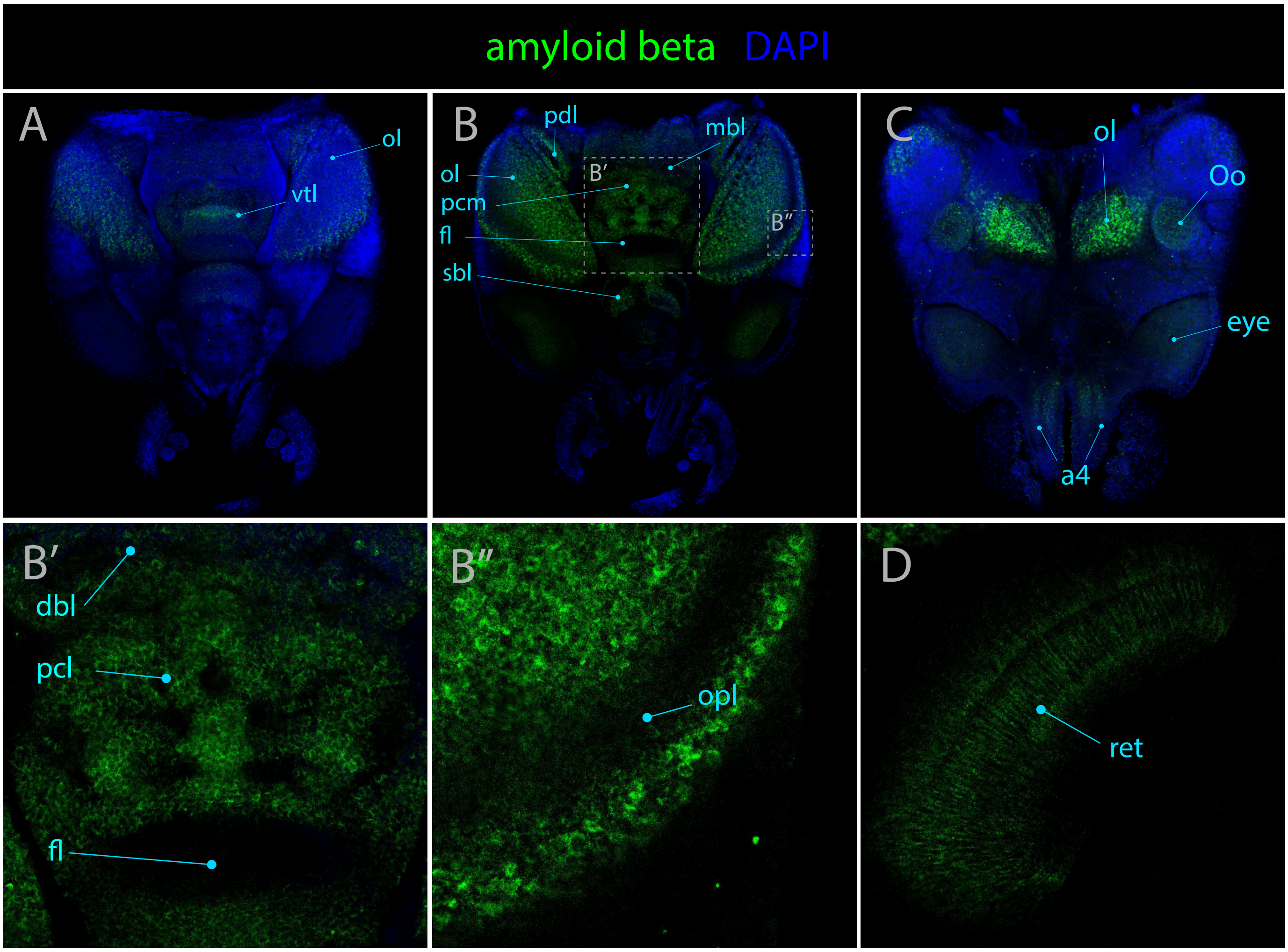
amyloid beta DAPI
A
ol
vtl
B
pdl
mbl
ol
B′
pcm
fl
B″
sbl
C
ol
Oo
eye
a4
B′
dbl
pcl
fl
B″
opl
D
ret

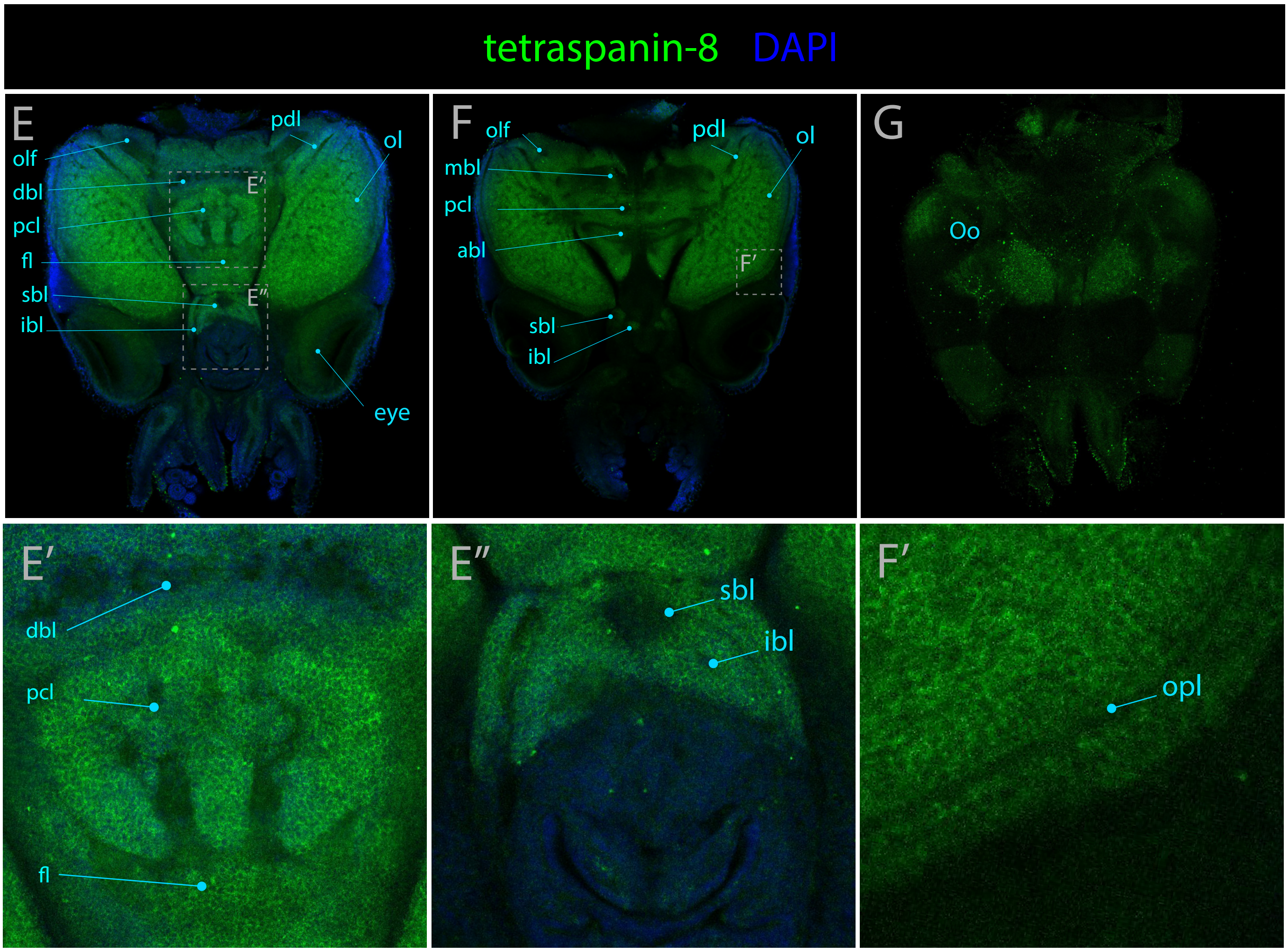
tetraspanin-8 DAPI
E
pdl
olf
ol
dbl
E′
pcl
fl
sbl
E″
ibl
eye
F
olf
pdl
ol
mbl
pcl
abl
F′
sbl
ibl
G
Oo
E′
dbl
pcl
fl
E″
sbl
ibl
F′
opl

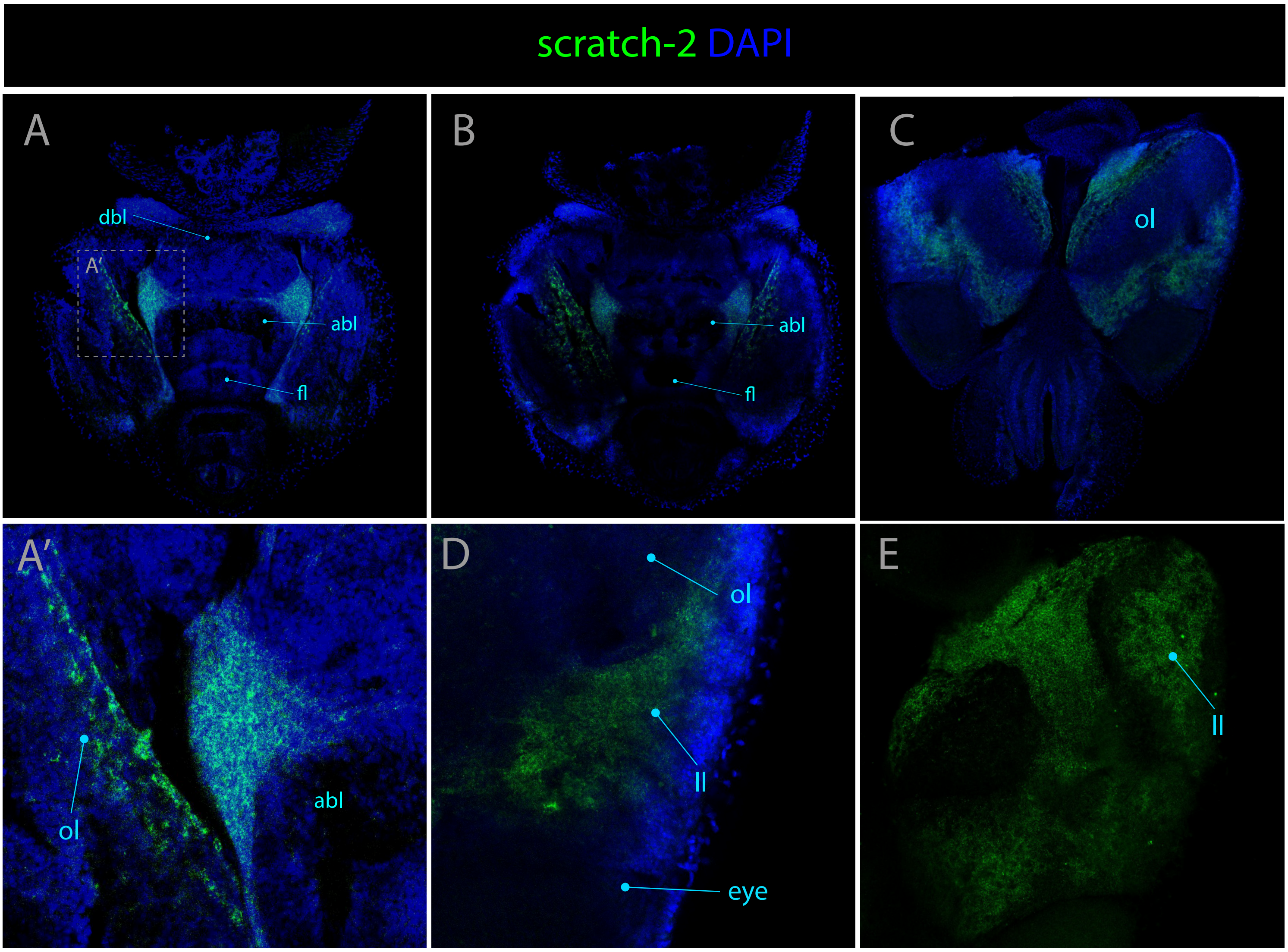

scratch-2 DAPI
A
dbl
A′
abl
fl
B
abl
fl
C
ol
A′
ol
abl
D
ol
ll
eye
E
ll

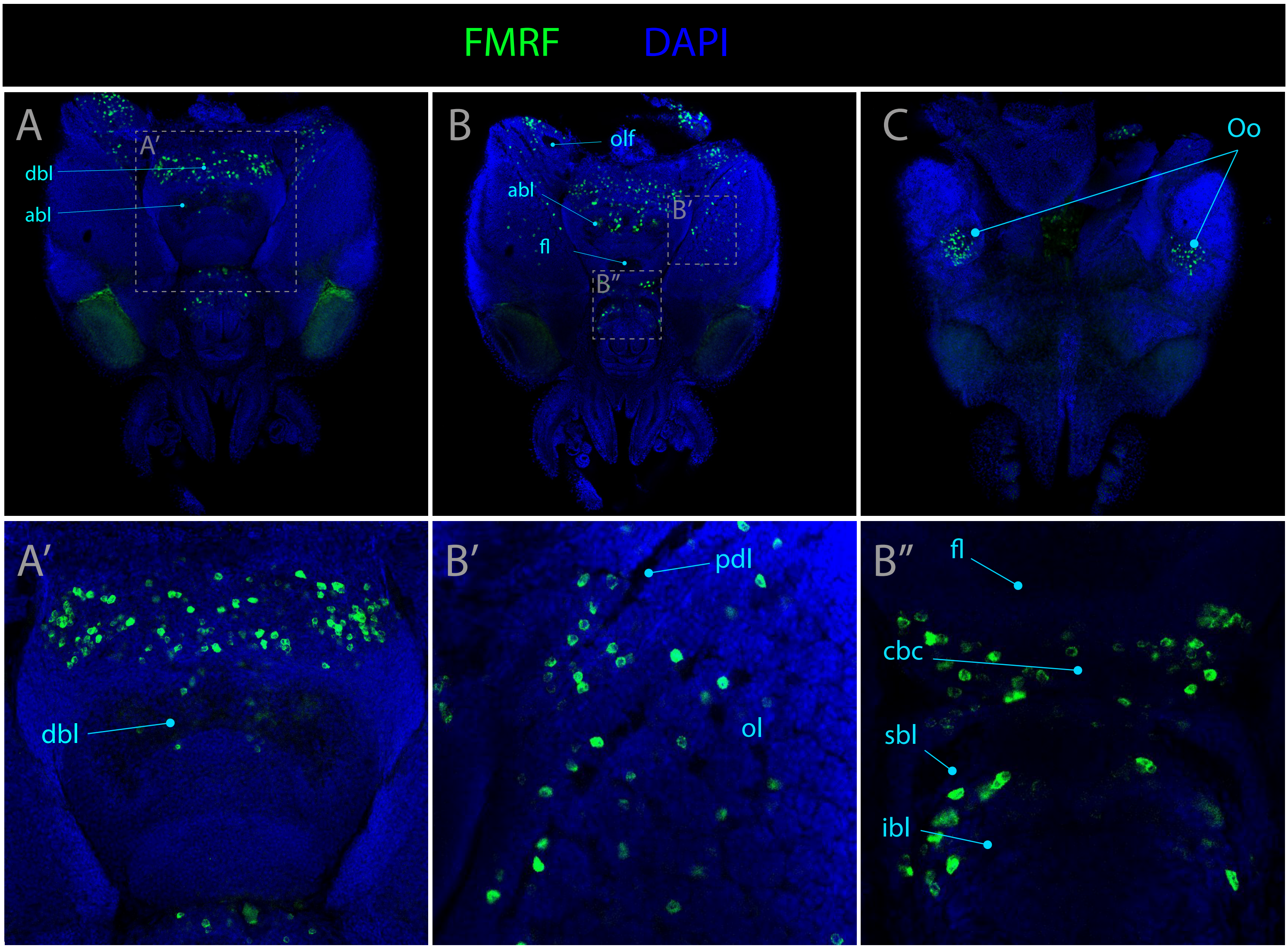
FMRF
DAPI
A
A'
dbl
abl
B
olf
abl
fl
B'
B''
C
Oo
A'
dbl
B'
pdl
ol
B''
fl
cbc
sbl
ibl

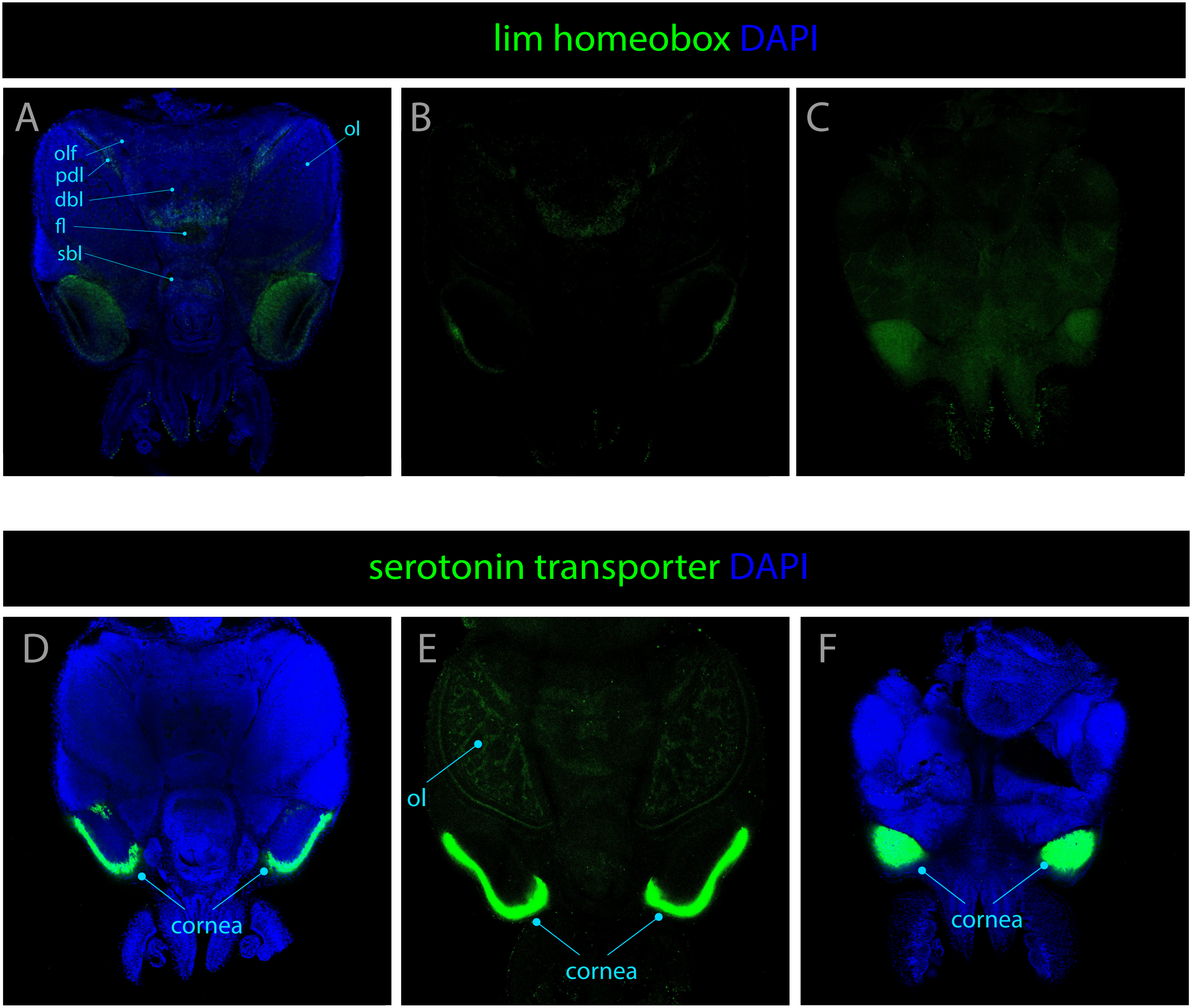
lim homeobox DAPI
A
olf
pdl
dbl
fl
sbl
ol
B
C
serotonin transporter DAPI
D
cornea
E
ol
cornea
F
cornea

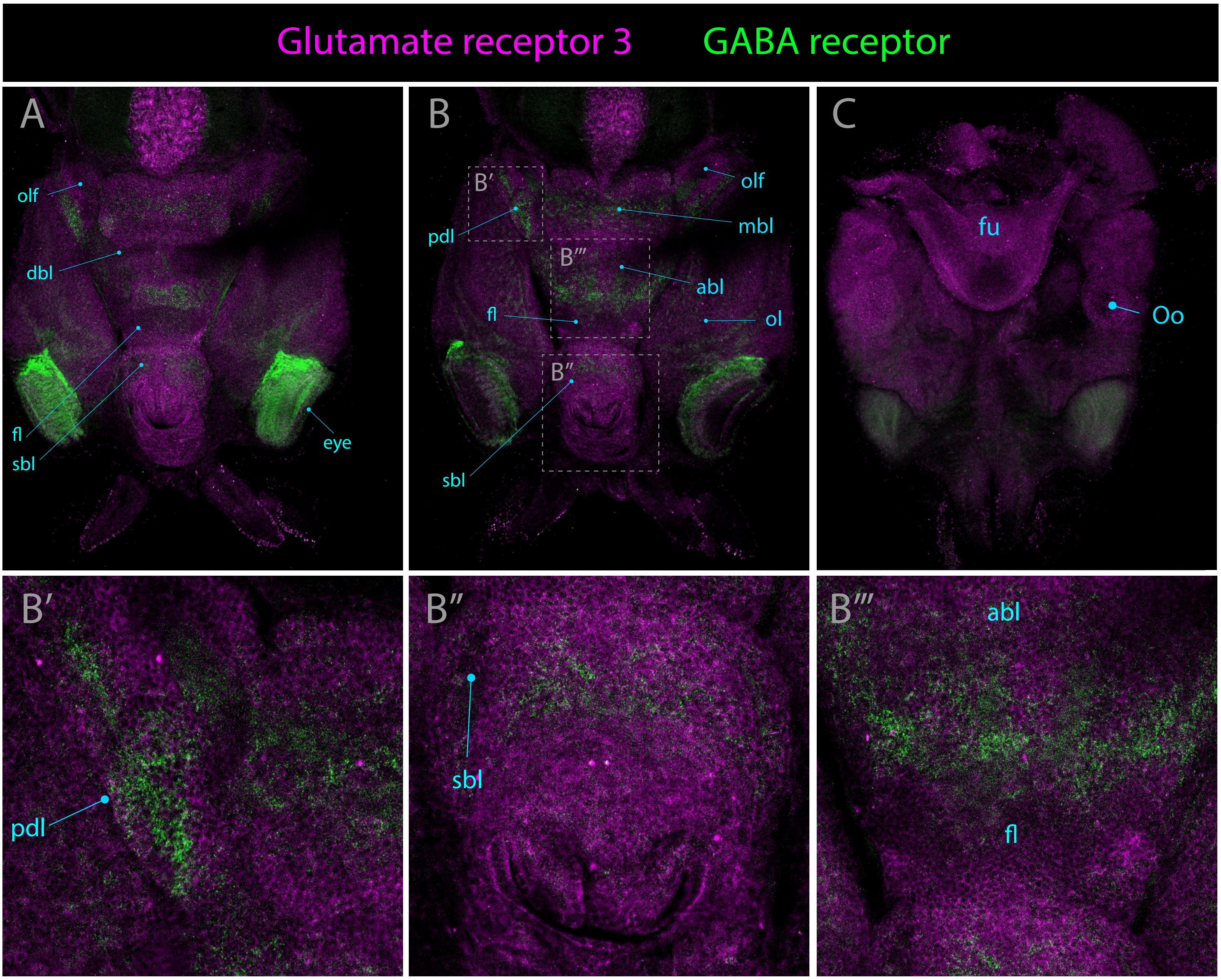
Glutamate receptor 3
GABA receptor
A
olf
dbl
fl
sbl
eye
B
B′
pdl
olf
mbl
B‴
abl
fl
ol
B″
sbl
C
fu
Oo
B′
pdl
B″
sbl
B‴
abl
fl

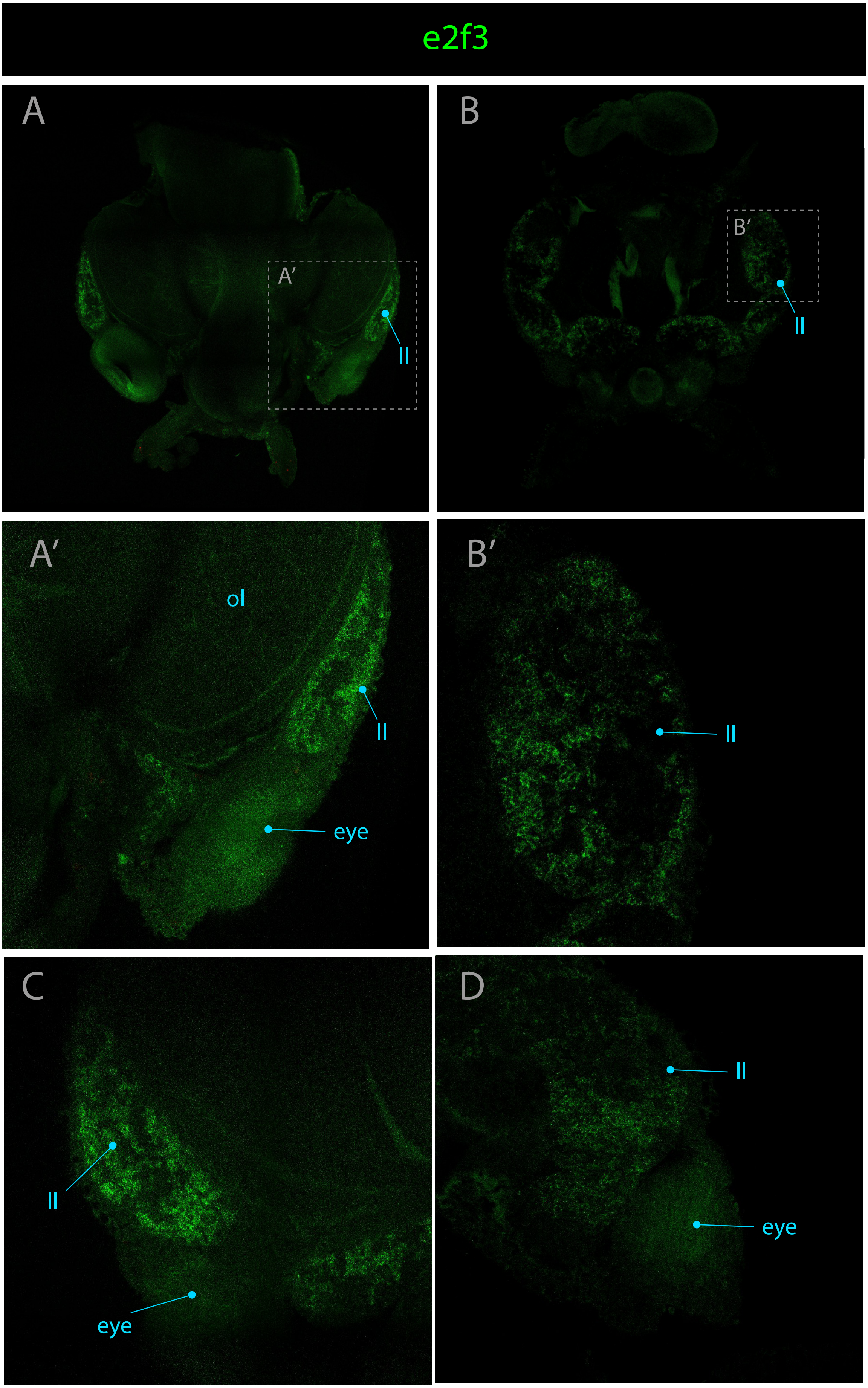
e2f3
A
A′
ll
B
B′
ll
A′
ol
ll
eye
B′
ll
C
ll
eye
D
ll
eye

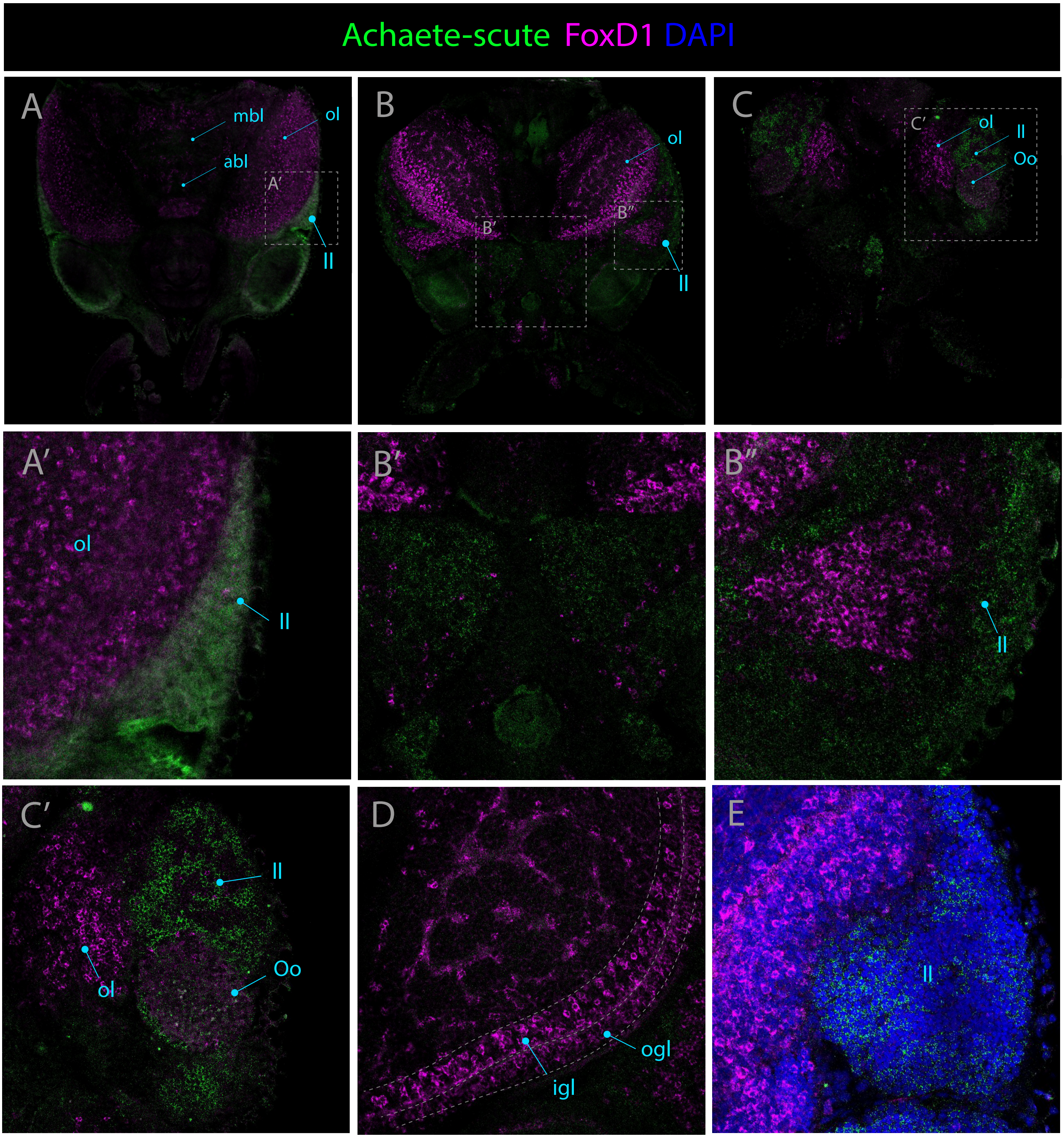
Achaete-scute FoxD1 DAPI
A
mbl
ol
abl
A′
ll
B
ol
B′
B″
ll
C
C′
ol
ll
Oo
A′
ol
ll
B′
B″
ll
C′
ll
ol
Oo
D
igl
ogl
E
ll

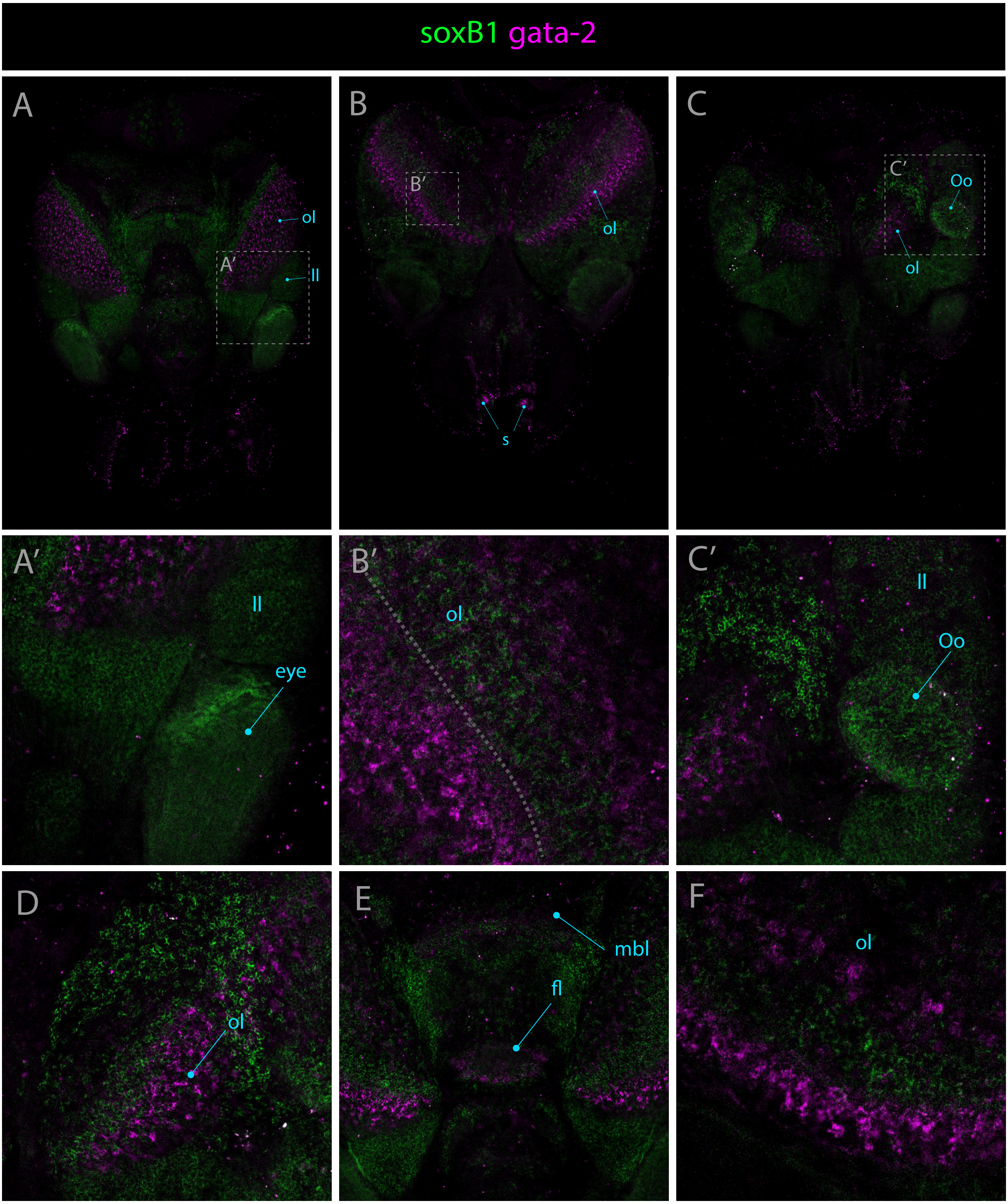

soxB1 gata-2
A
ol
A′
ll
B
B′
ol
s
C
C′
Oo
ol
A′
ll
eye
B′
ol
C′
ll
Oo
D
ol
E
mbl
fl
F
ol

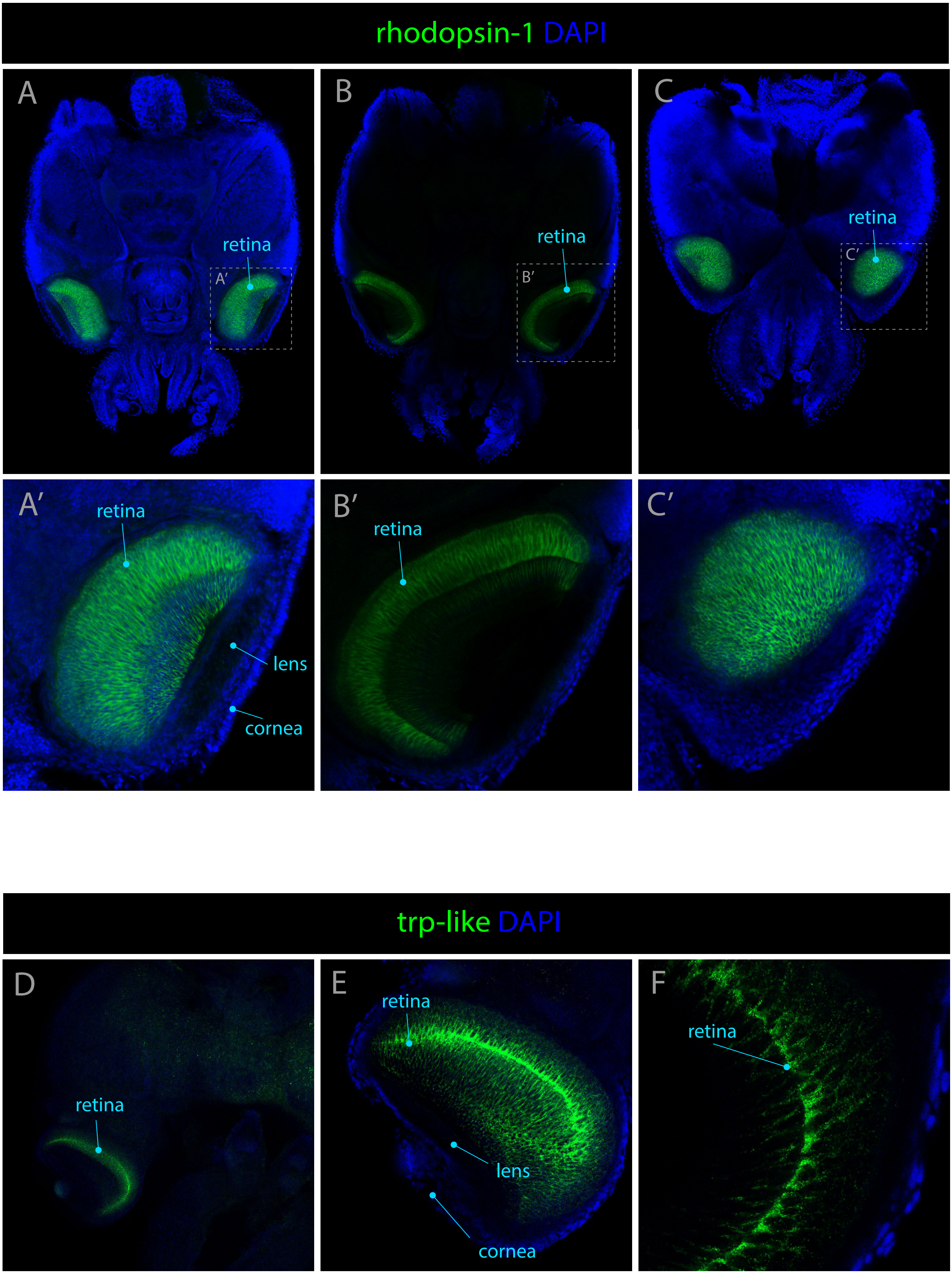

rhodopsin-1 DAPI
A
retina
A′
B
retina
B′
C
retina
C′
A′
retina
lens
cornea
B′
retina
C′
trp-like DAPI
D
retina
E
retina
lens
cornea
F
retina

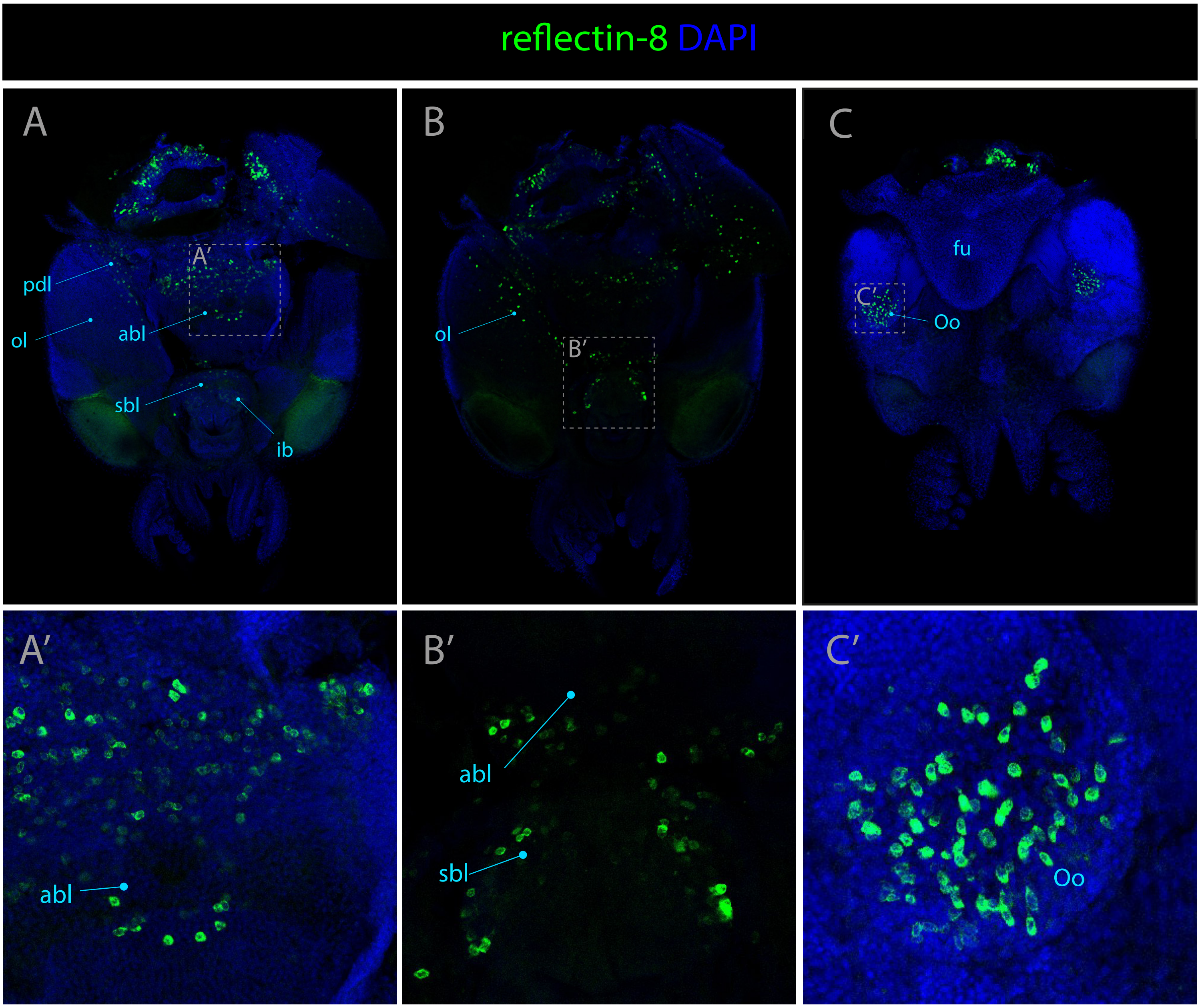

reflectin-8 DAPI
A
A′
pdl
ol
abl
sbl
ib
B
B′
ol
C
fu
C′
Oo
A′
abl
B′
abl
sbl
C′
Oo

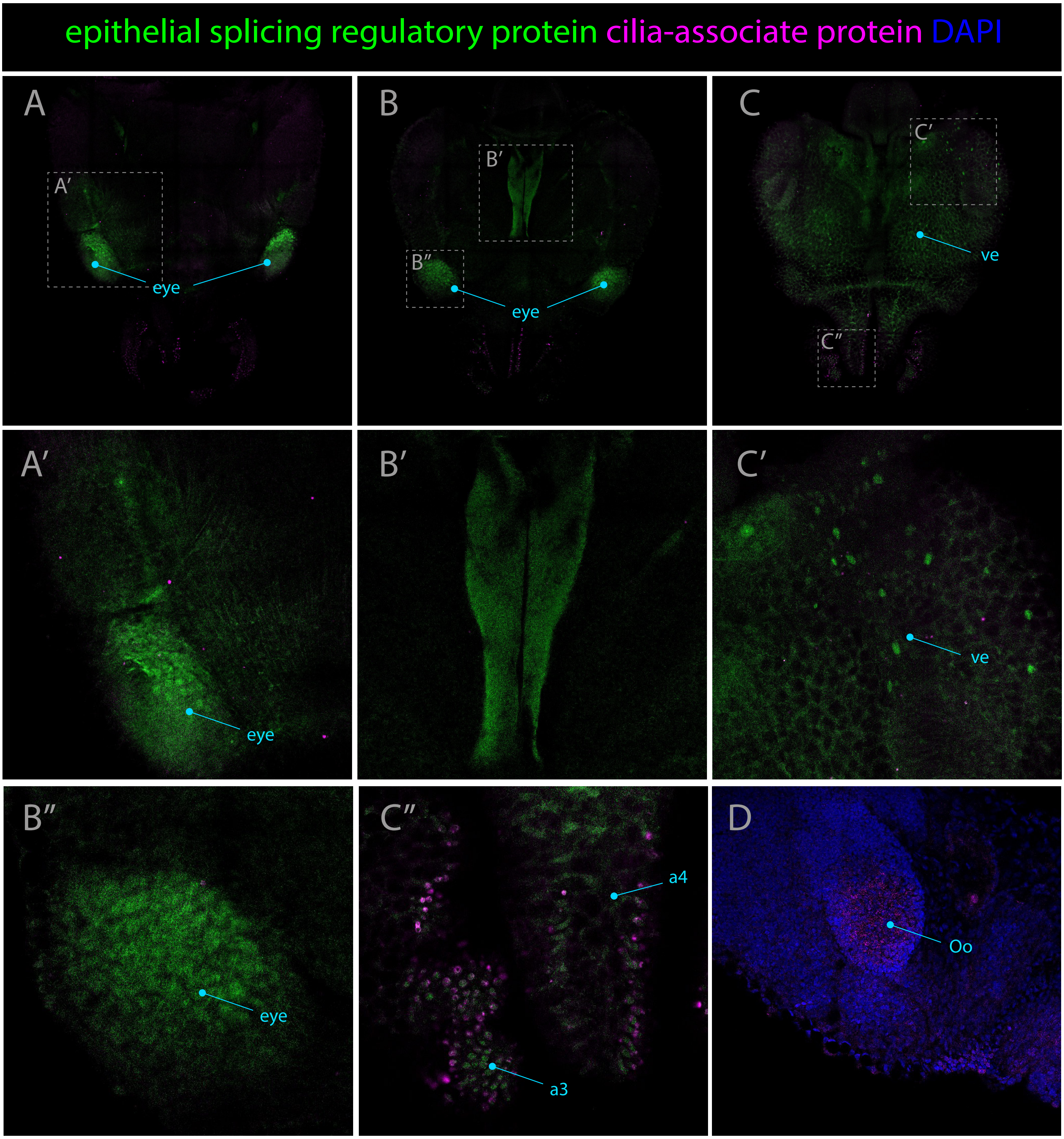

epithelial splicing regulatory protein cilia-associate protein DAPI
A
A′
eye
B
B′
B″
eye
C
C′
C″
ve
A′
eye
B′
C′
ve
B″
eye
C″
a4
a3
D
Oo

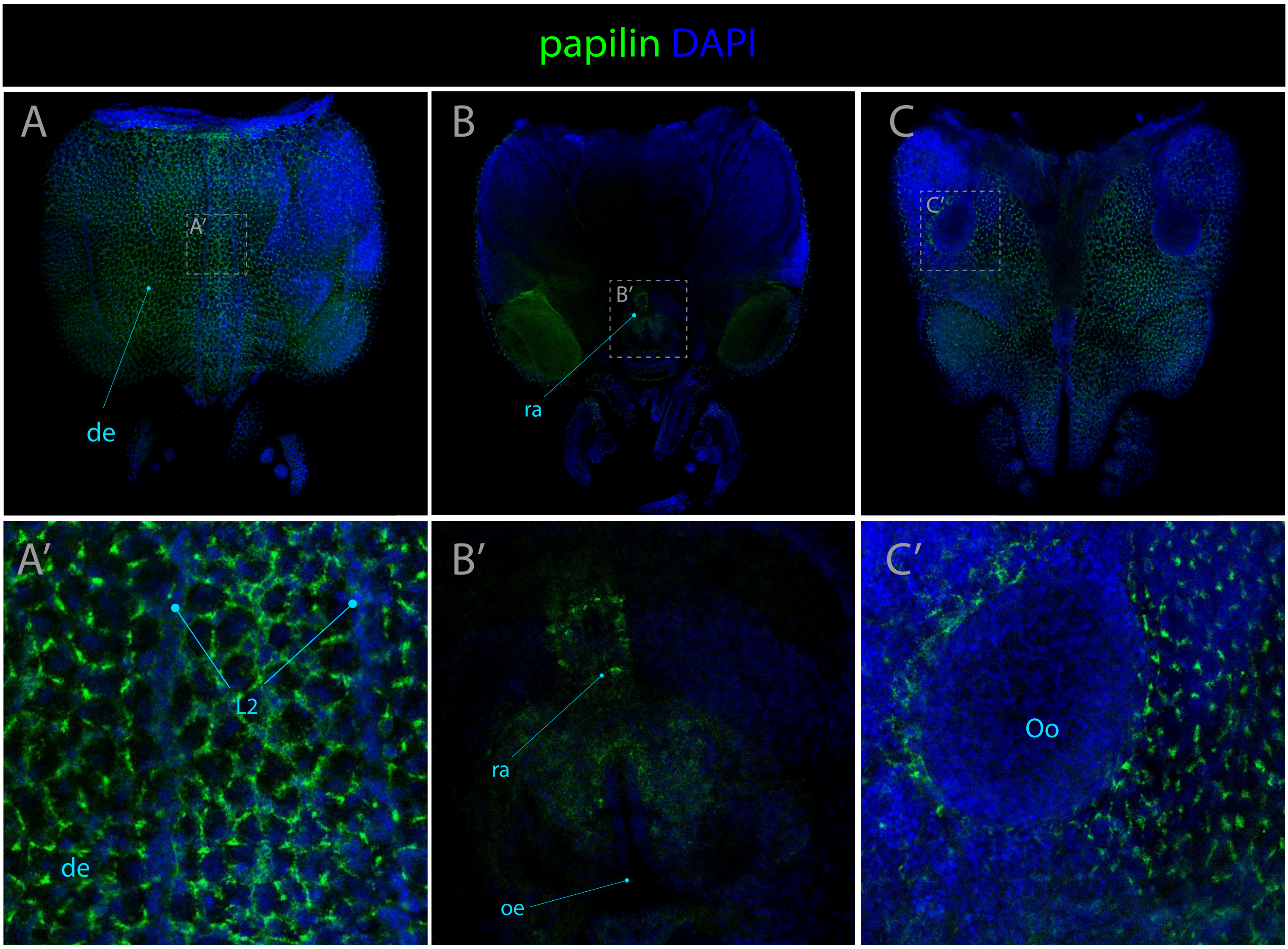
papilin DAPI
A
B
C
A'
B'
C'
de
ra
L2
oe
Oo

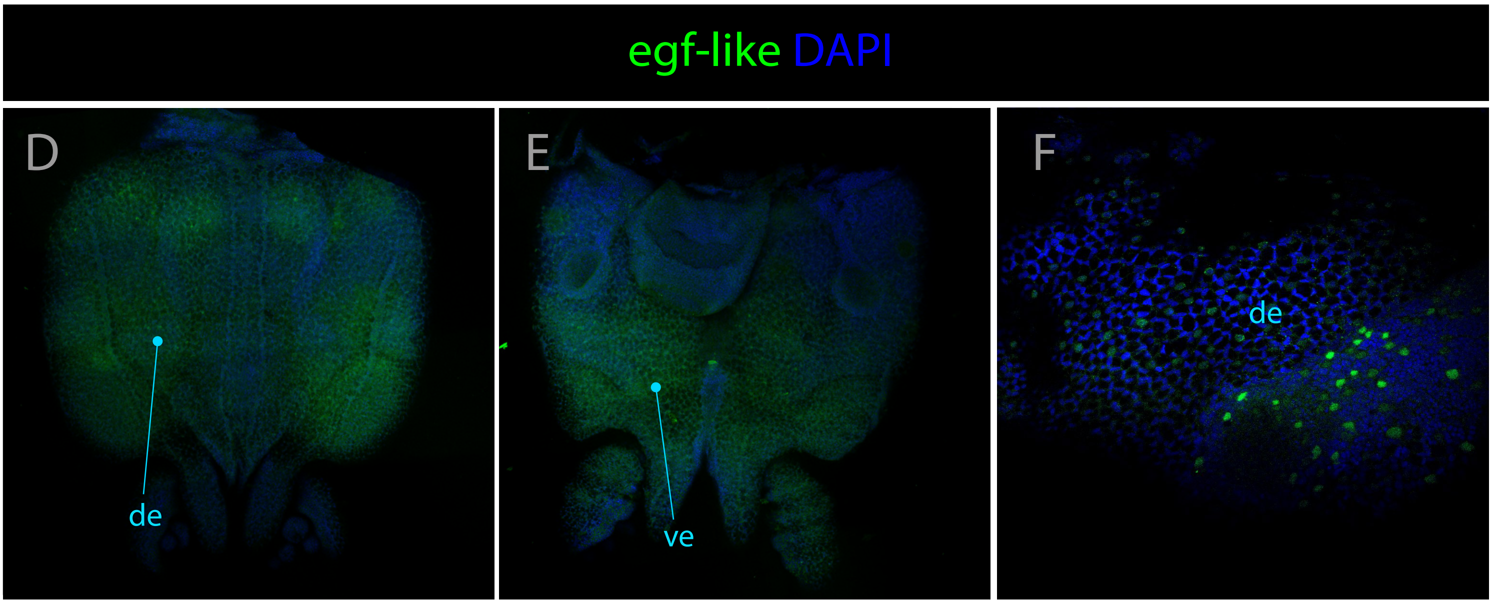
egf-like DAPI
D
E
F
de
ve

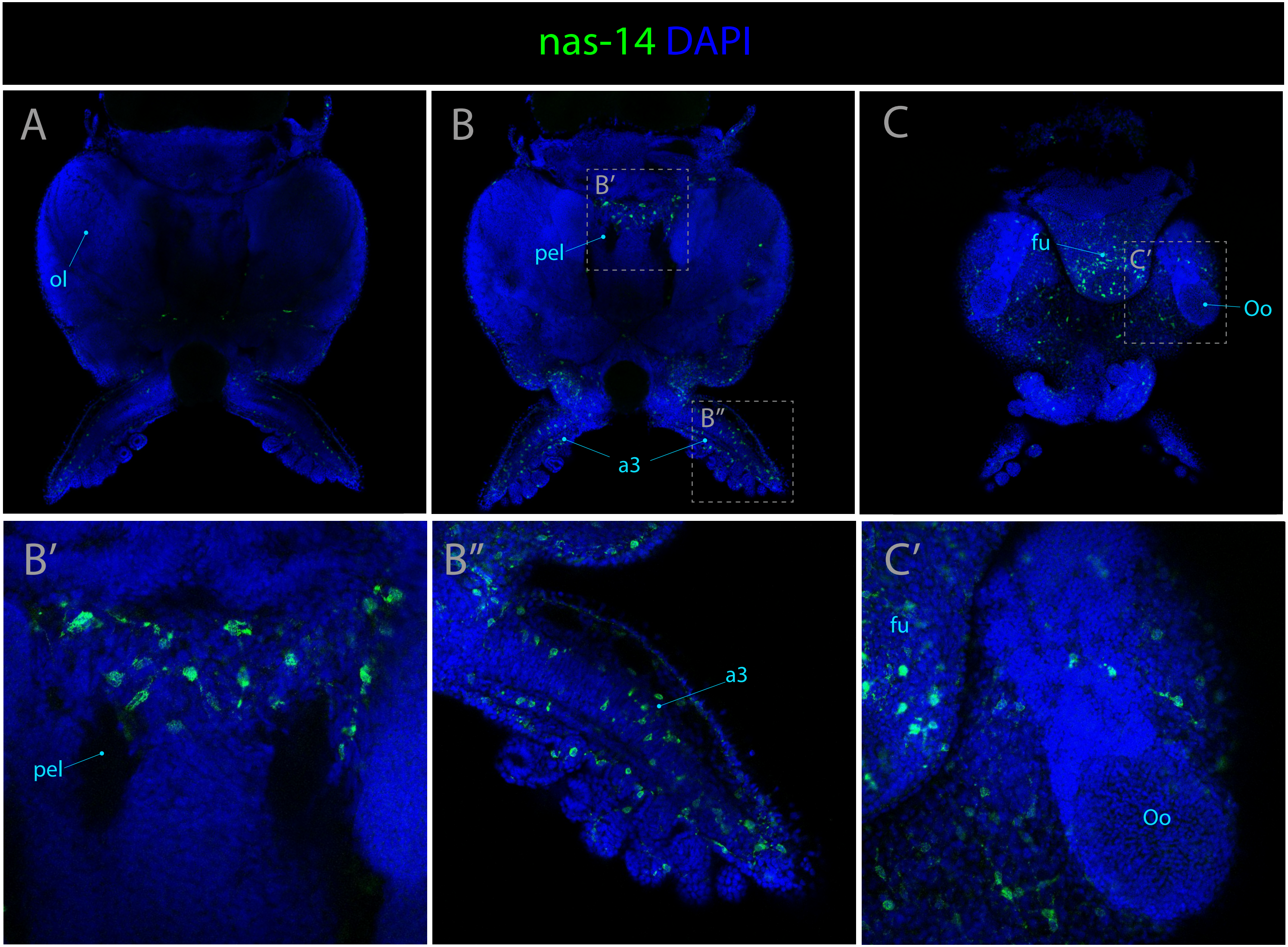
nas-14 DAPI
A
ol
B
B′
pel
B″
a3
C
fu
C′
Oo
B′
pel
B″
a3
C′
fu
Oo

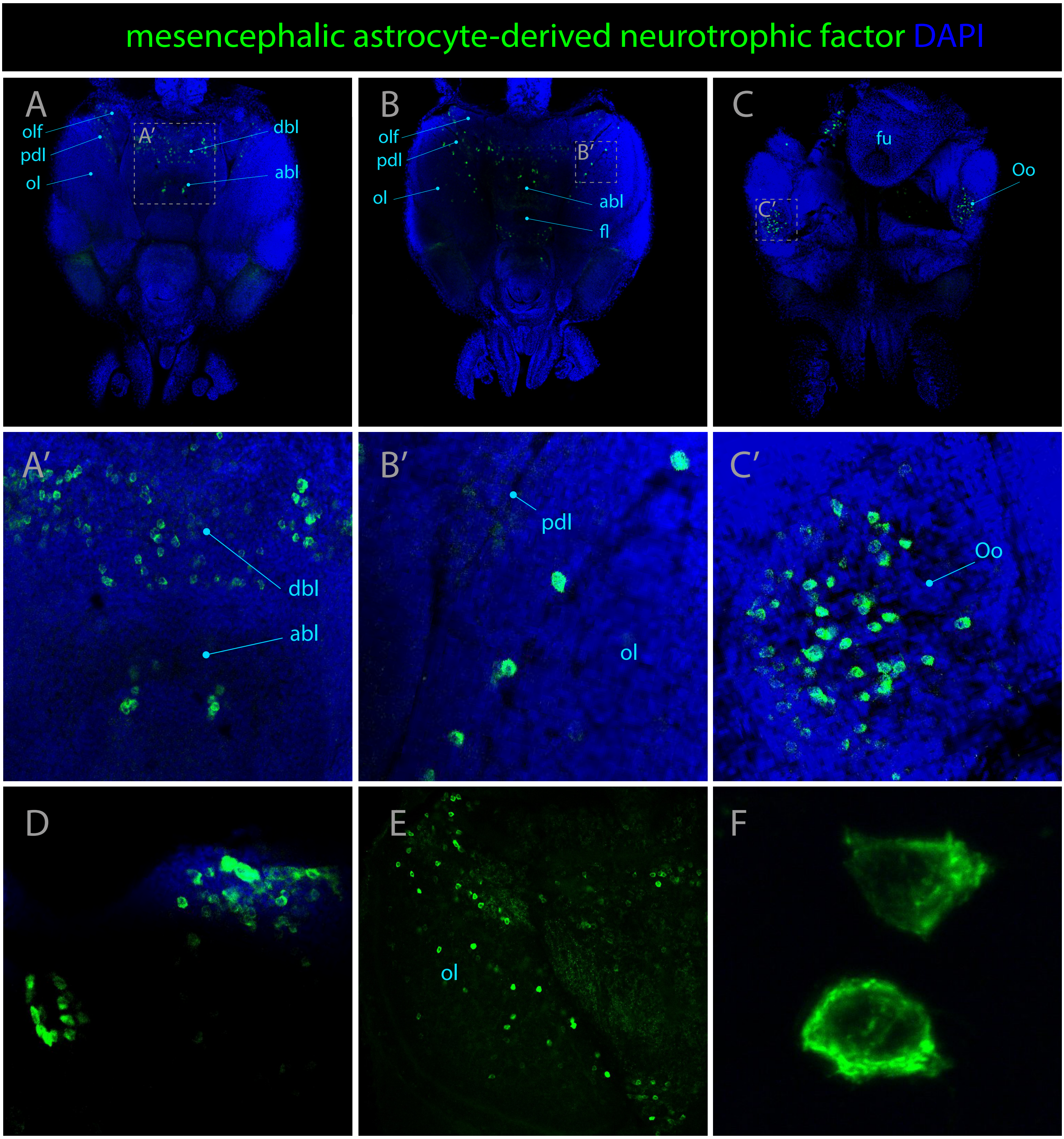
mesencephalic astrocyte-derived neurotrophic factor DAPI
A
olf
pdl
ol
A′
dbl
abl
B
olf
pdl
ol
B′
abl
fl
C
fu
Oo
C′
A′
dbl
abl
B′
pdl
ol
C′
Oo
D
E
ol
F

**Supplementary figure 1 : Expression patterns of *elav-like* and *cugpb-elav***

Confocal micrographs showing hybridization chain reactions for *elav-like* (green) and *cugpb-elav* (magenta). The areas shown are: (**A/A'/A''**) Dorsal overview **(B)** Medial overview **(C/C')** ventral overview **(D)** Dorsal view around the eye region **(E)** dorsal epidermis **(F)** arm **(G/G')** brain, circled area highlights an area of differential expression between the two genes. Abbreviations: (a3/a4) arm pairs 3 and 4, (abl) anterior basal lobe, (bm) buccal mass, (fl) frontal lobe, (fu) funnel, (ll) lateral lip, (ibl) inferior buccal mass, (ipl) inner plexiform layer , (L1) epidermal lines 1, (mbl) medial basal lobe, (ol) optic lobe, (Oo) olfactory organ, (opl) outer plexiform layer, (pcl) precommissural lobe, (ret) retina, (sbl) superior buccal lobe.

**Supplementary figure 2 : Expression patterns of *amyloid-beta* and *tetraspanin-8***

Confocal micrographs showing hybridization chain reactions for *amyloid-beta* and *tetraspanin-8*, co-stained with DAPI (blue). The areas shown are: (**A**) Dorsal overview, **(B/B'/B'')** Medial overview, **(C)** ventral overview, **(D)** eye, **(E/E'/E'')** dorsal overview, **(F/F')** medial overview, **(G)** ventral overview. Abbreviations: (a4) arm pair 4, (abl) anterior basal lobe, (bm) buccal mass, (dbl) dorsal basal lobe, (fl) frontal lobe, (fu) funnel, (ll) lateral lip, (ibl) inferior buccal mass, (ipl) inner plexiform layer , (L1) epidermal lines 1, (mbl) medial basal lobe, (ol) optic lobe, (olf) olfactory lobe, (Oo) olfactory organ, (opl) outer plexiform layer, (pdl) peduncle lobe, (pcl) precommissural lobe, (ret) retina, (sbl) superior buccal lobe, (vtl) vertical lobe.

**Supplementary figure 3 : Expression pattern of *scratch-2***

Confocal micrographs showing hybridization chain reactions for *scratch-2* (green), co-stained with DAPI (blue). The areas shown are: (**A/A'**) Dorsal overview, **(B)** Medial overview, **(C)** ventral overview, **(D)** dorsal view of the lateral lip above the eye region, **(E)** ventral view around optic lobe without DAPI. Abbreviations: (abl) anterior basal lobe, (dbl) dorsal basal lobe, (fl) frontal lobe, (ll) lateral lip, (ol) optic lobe, (olf) olfactory lobe.

**Supplementary figure 4 : Expression pattern of *fmrf***

Confocal micrographs showing hybridization chain reactions for *fmrf* (green), co-stained with DAPI (blue). The areas shown are: (**A/A'**) Dorsal overview, **(B/B'/B'')** Medial overview, **(C)** ventral overview. Abbreviations: (abl) anterior basal lobe, (dbl) dorsal basal lobe, (cbc) cerebro-buccal connective, (fl) frontal lobe, (ibl) inferior buccal mass, (ol) optic lobe, (olf) olfactory lobe, (Oo) olfactory organ, (pdl) peduncle lobe, (sbl) superior buccal lobe.

**Supplementary figure 5 : Expression patterns of *lim homeobox* and *serotonin transporter***

Confocal micrographs showing hybridization chain reactions for *lim homeobox* (green) and *serotonin transporter* (green), co-stained with DAPI (blue). The areas shown are: (**A**) Dorsal overview, **(B)** Medial overview, **(C)** ventral overview, **(D)** dorsal overview, **(E)** medial overview, **(F)** ventral overview. Abbreviations:
(dbl) dorsal basal lobe, (fl) frontal lobe, (ol) optic lobe, (olf) olfactory lobe, (pdl) peduncle lobe, (sbl) superior buccal lobe.

**Supplementary figure 6 : Expression patterns of *glutamate receptor 3* and *gaba receptor***

Confocal micrographs showing hybridization chain reactions for *glutamate receptor 3* (magenta) and *gaba receptor* (green), co-stained with DAPI (blue). The areas shown are: (**A**) Dorsal overview, **(B/B'/B'')** Medial overview, **(C)** ventral overview. Abbreviations: (abl) anterior basal lobe, (dbl) dorsal basal lobe, (fl) frontal lobe, (fu) funnel, (ll) lateral lip, (mbl) medial basal lobe, (ol) optic lobe, (olf) olfactory lobe, (Oo) olfactory organ, (pdl) peduncle lobe, (sbl) superior buccal lobe.

**Supplementary figure 7 : Expression pattern of *e2f3***

Confocal micrographs showing hybridization chain reactions for *e2f3* (green). The areas shown are: (**A/A'**) Dorsal overview, **(B/B')** ventral overview **(C)** dorsal part of the lateral lip around the eye **(D)** ventral part of the lateral lip around the eye. Abbreviations: (ll) lateral lip, (ol) optic lobe.

**Supplementary figure 8 : Expression patterns of *achaete-scute* and *foxD1***

Confocal micrographs showing hybridization chain reactions for *achaete-scute* (green) and *foxD1* (magenta), co-stained with DAPI (blue). The areas shown are: (**A/A'**) Dorsal overview, **(B/B'/B'')** Medial overview, **(C/C')** ventral overview, **(D)** optic lobe **(E)** ventral view of the lateral lip. Abbreviations: (abl) anterior basal lobe, (igl) inner granular layer, (ll) lateral lip, (mbl) medial basal lobe, (ol) optic lobe, (ogl) outer granular layer, (Oo) olfactory organ.

**Supplementary figure 9 : Expression patterns of *soxB1* and *gata-2***

Confocal micrographs showing hybridization chain reactions for *soxB1* (green) and *gata-2* (magenta), co-stained with DAPI (blue). The areas shown are: (**A/A'**) Dorsal overview, **(B/B')** Medial overview, **(C/C')** ventral overview, **(D)** ventral extremity of the optic lobe, **(E)** dorsal view of the brain, **(F)** anterior part of the optic lobe. Abbreviations: (fl) frontal lobe, (ll) lateral lip, (mbl) medial basal lobe, (ol) optic lobe, (Oo) olfactory organ, (s) suckers.

**Supplementary figure 10 : Expression patterns of *rhodopsin-1* and *trp-like***

Confocal micrographs showing hybridization chain reactions for *rhodopsin-1* and *trp-like*, co-stained with DAPI (blue). The areas shown are: (**A/A'**) Dorsal overview, **(B/B')** Medial overview, **(C/C')** ventral overview, **(D)** Dorsal overview, **(E)** eye, **(F)** retina.

**Supplementary figure 11 : Expression pattern of *reflectin-8***

Confocal micrographs showing hybridization chain reactions for *reflectin-8* (green), co-stained with DAPI (blue). The areas shown are: (**A/A'**) Dorsal overview, **(B/B')** Medial overview, **(C/C')** ventral overview. Abbreviations: (abl) anterior basal lobe, (fu) funnel, (ibl) inferior buccal lobe, (ol) optic lobe, (olf) olfactory lobe, (Oo) olfactory organ, (pdl) peduncle lobe, (sbl) superior buccal lobe.

**Supplementary figure 12 : Expression patterns of the *epithelial splicing regulatory protein* and *cilia-associated protein* genes**

Confocal micrographs showing hybridization chain reactions for *epithelial splicing regulatory protein* (green) and *cilia-associated protein* (magenta). The areas shown are: (**A/A'**) Dorsal overview, **(B/B'/B")** Medial overview, **(C/C'/C")** ventral overview, **(D)** ventral surface around the olfactory organ, co-stained with DAPI (blue). Abbreviations: (a3/a4) arm pairs 3 and 4, (Oo) olfactory organ, (ve) ventral epidermis.

**Supplementary figure 13 : Expression patterns of *papilin* and *egf-like***

Confocal micrographs showing hybridization chain reactions for *papilin* and *egf-like*, co-stained with DAPI (blue). The areas shown are: (**A/A'**) Dorsal overview, **(B/B')** Medial overview, **(C/C')** ventral overview, (**D**) Dorsal overview, **(E)** Medial overview, **(F)** ventral overview. Abbreviations: (de) dorsal epidermis, (L2)epidermal lines 2 (oe) oesophagus, (Oo) olfactory organ, (ra) radula, (ve) ventral epidermis.

**Supplementary figure 14 : Expression patterns of *nas-14***

Confocal micrographs showing hybridization chain reactions for *nas-14,* co-stained with DAPI (blue). The areas shown are: (**A**) Dorsal overview, **(B/B'/B")** Medial overview, **(C/C')** ventral overview. Abbreviations: (a3) arm pair number 3, (fu) funnel, (oe) oesophagus, (ol) optic lobe, (Oo) olfactory organ, (pel) pedal lobe.

**Supplementary figure 15 : Expression pattern of *mesencephalic astrocyte derived neurotrophic factor***

Confocal micrographs showing hybridization chain reactions for *mesencephalic astrocyte derived neurotrophic factor* (green), co-stained with DAPI (blue). The areas shown are: (**A/A'**) Dorsal overview, **(B/B')** Medial overview, **(C/C')** ventral overview, (D) region adjacent to the funnel (E) optic lobe without DAPI staining, (F) cells at high magnification. Abbreviations: (abl) anterior basal lobe, (dbl) dorsal basal lobe, (fl) frontal lobe, (fu) funnel, (ol) optic lobe, (olf) olfactory lobe, (Oo) olfactory organ, (opl) outer plexiform layer, (pdl) peduncle lobe.
